# An expanded realm of anti-CRISPR-associated proteins and regulatory mechanisms

**DOI:** 10.64898/2026.07.27.740549

**Authors:** Maximilian Feussner, Nils Birkholz, So Yeon Lee, Chase L. Beisel, Hyun Ho Park, Chris M. Brown, Peter C. Fineran, Zasha Weinberg

**Affiliations:** Bioinformatics Group, Department of Computer Science and Interdisciplinary Centre for Bioinformatics, Leipzig University, 04107 Leipzig, Germany; Botnar Institute of Immune Engineering, 4056 Basel, Switzerland; Department of Microbiology and Immunology, University of Otago, PO Box 56, Dunedin, New Zealand; Bioprotection Aotearoa, University of Otago, PO Box 56, Dunedin, New Zealand; Genetics Otago, University of Otago, PO Box 56, Dunedin, New Zealand; Maurice Wilkins Centre for Molecular Biodiscovery, University of Otago, PO Box 56, Dunedin, New Zealand; College of Pharmacy, Chung-Ang University, Seoul 06974, Republic of Korea; Department of Global Innovative Drugs, Graduate School of Chung-Ang University, Seoul 06974, Republic of Korea; Department of Biochemistry, University of Otago, PO Box 56, Dunedin, New Zealand

**Author notes:** Correspondence may also be addressed to Peter C. Fineran. Institute of Computer Science and Institute of Biochemistry and Biotechnology, Martin-Luther University Halle-Wittenberg, 06114 Halle, Germany (Peter C. Fineran, Zasha Weinberg). Equal contribution (Maximilian Feussner, Nils Birkholz).

## Abstract

Many bacteriophages encode anti-CRISPR (Acr) proteins that inhibit the CRISPR-Cas immune systems. Rapid *acr* gene expression upon phage entry enables CRISPR-Cas neutralisation, but can impact phage fitness if unregulated. Therefore, Acr production is often controlled by distinct families of co-encoded anti-CRISPR-associated (Aca) proteins, which are usually helix-turn-helix (HTH) regulators that bind DNA within *acr*–*aca* operon promoters. Previously, we demonstrated that the Aca2 family additionally represses Acr production translationally by binding structured RNA motifs within the 5′ untranslated regions (UTRs) of the *acr–aca* mRNA. Here, through systematic bioinformatic analyses, we provide evidence of structured RNA motifs in the 5′ UTRs of operons encoding members of other Aca families, and that Aca1 also specifically binds its cognate RNA motif. Additionally, many Aca proteins are predicted to regulate not only their own but also adjacent operons with potential anti-defence genes. Indeed, we show that Aca14, newly identified in this study, represses two predicted anti-defence operons. Aca14 is a ribbon-helix-helix domain protein, revealing regulatory diversity beyond the canonical HTH Aca family members. Collectively, our findings expand the understanding of *acr* regulation in mobile genetic elements and reveal novel mechanisms by which phages fine-tune anti-defence gene expression.

**Graphical abstract:** 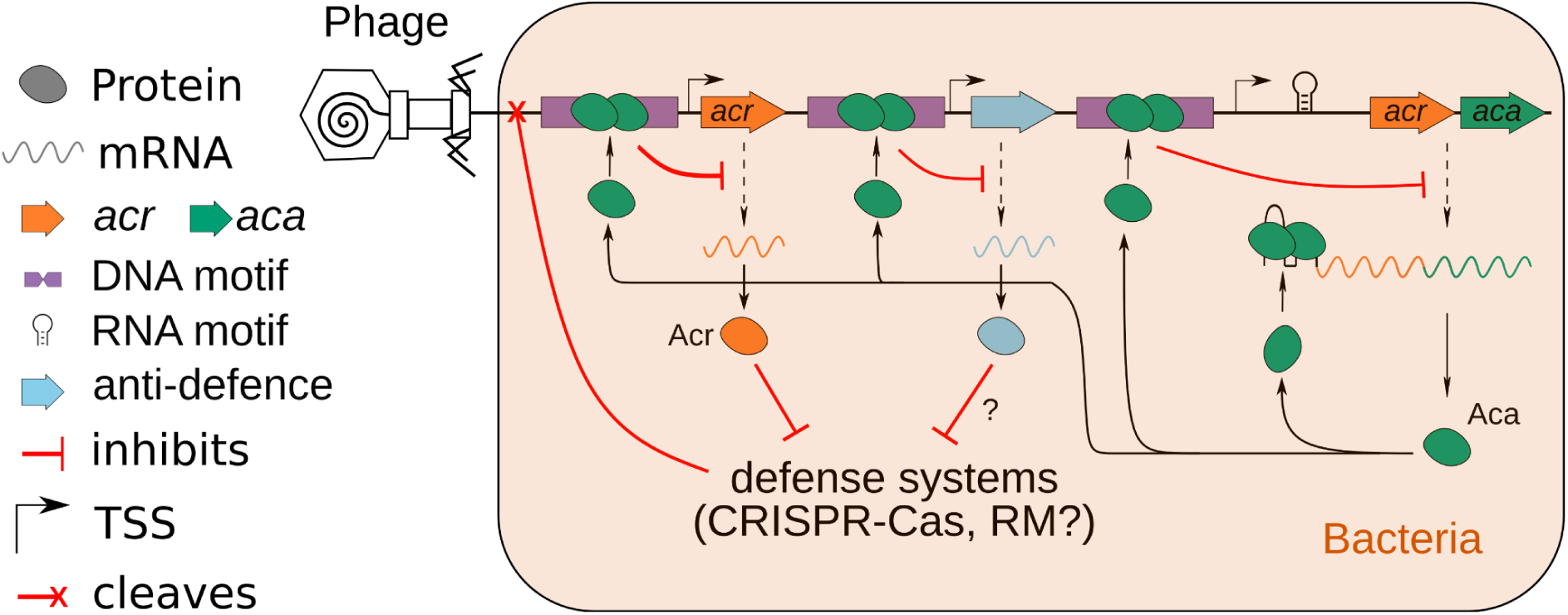

## Introduction

Bacteriophages (phages) are viruses that rely on their bacterial host for replication, and are responsible for 20-40% of daily bacterial turnover(1). To counteract the constant challenge from phage infection, bacteria have developed defence systems against phages, such as CRISPR-Cas and restriction-modification systems(2). However, phages have evolved various strategies to evade these defences during the evolutionary arms race with their bacterial hosts(3). A prominent example is the inhibition of CRISPR-Cas systems by phage-encoded anti-CRISPR proteins (Acrs) (4) or RNAs (Racrs) (5, 6). Acrs are highly diverse, with more than 100 examples known(7), and play an important part of the CRISPR-Cas biotechnological toolkit(8), such as regulating genome editing by Cas9 to reduce off-targeting. Although some Acrs function enzymatically, most bind CRISPR-Cas components to prevent targeting of foreign nucleic acids(9).

To inhibit CRISPR-Cas complexes, phages require rapid and robust production of Acr proteins immediately after infection. However, constitutive expression of Acr proteins can impair phage propagation by causing a metabolic or gene expression burden(10). Therefore, regulation of Acr expression is needed to enable inhibition of the host CRISPR-Cas systems, but then reduce Acr protein levels(11). Indeed, Acr proteins are regulated by Acr-associated (Aca) proteins that are frequently co-encoded in the same operon. Aca proteins form distinct families but generally contain a helix-turn-helix (HTH) domain, which is one of the most pervasive protein domains to facilitate gene regulation across all domains of life(12). The roles and mechanisms of the HTH domain are well-studied and were first described in 1981(13–15).

Bacteria use HTH proteins extensively to activate or repress gene expression. Canonically, these HTH regulators form dimers that bind inverted repeat DNA sequences by inserting one of the helices of the HTH motif into the major groove of DNA. If the binding sites in the DNA are located in or near promoters, HTH binding may inhibit transcription by preventing RNA polymerase (RNAP) access(12).

We previously demonstrated that Aca2 uses its HTH domain to bind not only the *acr-aca* promoter DNA and inhibit transcription(16, 17), but also to a widely conserved RNA motif in the 5′ UTR of its *acr-aca* operon. Aca2 binding to this structured RNA motif prevents Acr translation by blocking ribosome binding to the ribosomal binding site. Translational control by Aca2 is needed to control the expression of *acr* genes even at high DNA copy numbers that occur during phage replication. This Aca2-mediated regulatory mechanism appears to occur across diverse bacterial taxa, highlighting its broad relevance and expanding the conventional view that HTH domain proteins function predominantly as DNA-binding regulators(11).

Our previous discoveries raise broader questions: Do other *acr-aca* operons contain RNA motifs as potential binding sites of Aca proteins? Do other Aca proteins bind these motifs? And do Aca proteins have regulatory motifs for DNA and/or RNA binding in regions beyond *acr* operons? Here, to address these questions, we established a computational pipeline to systematically identify conserved motifs upstream of *acr-aca* operons. We identified a conserved RNA motif in the 5′ UTRs of *acr-aca1* operons and demonstrated binding of purified Aca1 to this motif. Comparative analysis of DNA- and RNA-binding motif locations across Aca families suggests that their regulatory roles often extend to neighbouring operons encoding Acr or other counter-defence proteins. Our experiments support this extended reach for a previously uncharacterized Aca protein containing a Ribbon-Helix-Helix (RHH) domain that we name Aca14, demonstrating that additional conserved protein domains mediate *acr* regulation. Collectively, our work underscores the broad regulatory potential of Aca proteins within bacterial defence networks.

## Material and Methods

### Pipeline to identify regulatory 5**′** UTRs associated with other Aca proteins

Operons encoding Aca1-12 were extracted from the IMG/VR database and treated as described previously(11, 17). In total, we extracted 4673 operons from 4628 different viral contigs. Aca13 was excluded because it lacked the conserved HTH domain that was, at the time of analyses, considered characteristic of verified Aca proteins. Given the small size and sequence variability of Aca proteins, we cannot exclude the possibility that some extracted operons may contain non-Aca HTH regulators(17). To identify potential RNA motifs, 100 nt upstream of the predicted start codon of *acr-aca* operons were used as an input for CMfinder(18). This 100 nt threshold was selected because previous studies have shown that the regulatory motifs of *acr-aca* operons predominantly reside within this region (11, 17, 19). The resulting potential RNA motifs were manually checked for covariance by comparing the number of covarying base pairs with the number of atypical basepairs relative to the consensus secondary structure. An alignment was considered promising only if the number of covarying base pairs exceeded the number of unpaired bases. Sequences were removed from promising alignments if the assigned CMfinder score was below 0.95 and if the predicted RNA secondary structure of the sequence had more than 2 unpaired bases compared to the consensus secondary structure. This strict threshold was applied to ensure structural stability, as even two unpaired bases can drastically alter the secondary conformation of small RNA molecules. Moreover, for each sequence, the nucleotides from the DNA motif to the start codon were extracted, and the resulting sequences were aligned with LocaRNA 2.0(20). This was done to include potential conserved ncRNA regions that were missed by CMfinder and to have all regulatory elements in one alignment. The output of LocaRNA was used as a seed alignment for a covariance model(21) (CM) and to search all operons from the respective Aca protein with an E-value cutoff 0.001. The same approach, including covariation search, was used to identify the DNA motif of the *acr-aca* operons. To validate these motifs and their positions, we searched for high-confidence transcription start sites (TSS) downstream of the motif exhibiting greater than 2000 transcriptional initiation rate units, predicted using the Promoter Calculator(22). Only DNA motifs supported by (i.e., located upstream of) such TSS predictions were considered high-confidence. For promising RNA motifs, the initial alignment output was used as a seed for a second search to identify more distant homologs. The results of that search were analysed with R-Scape (23) for statistically significant covariance and sequence conservation to identify conserved RNA motifs in the 5′ UTR. The RNA motif was considered conserved if it had at least two base pairs with statistically significant covariation (E-value < 0.05) and one base pair with highly conserved nucleotides (at least 97% conserved)(**Supplementary Figure S1**). For Aca6, Aca9, and Aca11, manual curation based on the previously described DNA motifs was required to identify the initial DNA motif. Because our pipeline was designed to detect conserved motifs within 100 nt upstream of acr–aca operons, it may under-detect families with more heterogeneous binding sites or more variable promoter architectures. The conserved DNA motifs were visualised with the MEME suite(24). The position of the RNA motifs relative to the TSS was predicted using the Promoter Calculator from the Salis Lab(22). This tool’s limitation to *Escherichia coli* data, combined with the diversity of Aca proteins, may explain why we were unable to identify DNA motifs with corresponding transcription start sites for all *acr-aca* operons. We surveyed public repositories for relevant experimental TSS datasets, but did not identify locus-resolved data applicable to the specific Aca-associated operons analysed here.

### Phylogenetic tree of the HTH domain of Aca1

In the first step, the operons containing only the Aca protein were removed from the non-redundant operon set because they did not have an RNA motif before their start codon. In a second step, the amino acid sequences of the Aca proteins were extracted from the operons using prokka_v (version 1.12-viral)(25). The predicted host of each operon was extracted from the IMG/VR database(26). The amino acid sequences were clustered at 90% (for Aca1) and aligned using T-COFFEE (https://www.ebi.ac.uk/jdispatcher/msa/tcoffee)(27). The resulting alignment was used as an input file with default parameters for FastTree (version 2.1.11)(28), and the tree was visualised using the web server Interactive Tree of Life(29).

### Clustering of RNA motifs to correlate with predicted hosts

To identify potential RNA subclusters in the Aca1 RNA motif, the RNA sequences matching the motif were clustered with CD-HIT(30) at a sequence identity threshold of 80%. Clusters with more than four sequences were visualised with the predicted host and the amino acid sequence of Aca1 via the phylogenetic tree using the web server Interactive Tree of Life(29). This analysis resulted in three RNA-clusters. To test whether RNA-cluster identity or host-family assignment was non-randomly distributed across the Aca1 phylogeny, we treated each annotation (one of three RNA-clusters or one of five bacterial families) as a discrete tip trait and calculated the Fitch parsimony score(31, 32) on the fixed tree. For the host-family analysis, only Aca1 proteins associated with one of the three RNA clusters were included. Significance was assessed by permutation testing, in which trait labels were randomized across tree tips and the parsimony score recalculated for each replicate (10, 000 times for Empirical one-sided p-value); significantly lower observed scores were interpreted as evidence of phylogenetic clustering. The analysis was repeated on unclustered, 95% identity-clustered, and 90% identity-clustered Aca1 datasets to assess robustness to sequence dereplication.

### Search for DNA and RNA motifs in the RefSeq database

The bacterial genomes from the RefSeq database (version 223)(33) were downloaded from the FTP server (https://ftp.ncbi.nlm.nih.gov/genomes/refseq). The downloaded genomes were searched with the CMs of Aca1 and Aca2 DNA and RNA motifs obtained from the above-described pipeline with the cmsearch program from the Infernal package(21) using an E-value cutoff of 0.1.

### Protein analysis of conserved domains and anti-defence function

All proteins encoded within 2000 nt up- and downstream of the predicted DNA- and RNA-binding motifs in the RefSeq bacterial genomes were downloaded via Biopython’s Entrez package(34). Proteins were predicted by prokka_v (version 1.12-viral)(25) for the operons of the IMG/VR database(26). Only genes that were presumably in the same operon (intergenic region <60 nt) as the gene directly downstream of the DNA- and RNA-binding site were used.

Conserved domains within the extracted proteins were searched using the web server of the Conserved Domain Database (https://www.ncbi.nlm.nih.gov/Structure/bwrpsb/bwrpsb.cgi)(35). Anti-defence functions were inferred using HMMs downloaded from the Anti-Prokaryotic Immune System Database (https://bcb.unl.edu/dbAPIS/index.php)(36) and the AntiDefenseFinder(37). The HMMs were used to scan the extracted proteins with the command ‘hmmscan’ from the HMMER suite(38), utilising the standard parameters. After the scan, we annotated proteins as Acr proteins if they matched the HMMs with an E-value lower than 10^-5^.

To cluster the proteins predicted using the above methods to be regulated by Aca proteins (excluding the Aca proteins themselves), a local BLAST database was constructed from these sequences using the BLAST+ suite. An all-against-all BLASTp search was then executed using default parameters. Protein sequences were grouped into clusters if they shared at least one pairwise alignment meeting the following criteria: an E-value < 10^-10^, a sequence identity > 40%, and a query coverage > 80%. Within each resulting cluster, the protein exhibiting the highest number of intra-cluster matches was designated as the representative sequence. Functional annotations were subsequently assigned to each cluster based on the majority consensus classification derived from AntiDefenseFinder or the NCBI Conserved Domain Database (CDD).

### Search for RHH-domain homologs and their DNA motifs

We used the first identified RHH-domain candidate Aca (WP_005725436.1) from *Pasteurella multocida* as input for PSI-BLAST via the BLAST webserver (accessed Sept. 6, 2025). After receiving the first results, we ran PSI-BLAST a second time to identify more distant homologs. We utilised our nucleotide motif discovery pipeline to identify the conserved DNA inverted repeats upstream of the *acr-rhh* operons. The conserved DNA motif was visualised with the MEME suite(24), and the position relative to the transcription starting site was predicted using the Promoter Calculator from the Salis Lab(22).

### Protein-DNA structure prediction of RHH protein with AlphaFold 3

The predictions were made via the AlphaFold 3 web server (https://alphafoldserver.com/, Accessed 25.06.2026) by submitting the amino acid sequence of the RHH protein from *Pasteurella multocida* and the predicted DNA motif associated with this protein. To test different stoichiometries of the RHH protein, simulations with one, two and four proteins, as well as two copies of the predicted DNA motif, were analysed(39). The results with the best pIDDT, ipTM, and pTM were chosen and downloaded (**Supplementary Table S2**). For visualisation, the model was loaded into PyMOL(40) (Aca1) or Mol* (RHH)(41). For the structural prediction of Aca1-binding DNA and RNA, the nucleotide motifs and amino acids were submitted to the AlphaFold 3 webserver (https://alphafoldserver.com/, Accessed 25.06.2026)(**Supplementary Table S3**).

### Purification of Aca1, Aca12 and Aca14

The *aca1* gene from *Pseudomonas aeruginosa* phage JBD30 (accession code: WP_016068277.1) and the *aca12* gene from bacteriophage sp. (accession code: DAX43943.1) were synthesised by Bionics (Daejeon, Republic of Korea). The *aca14* gene from *Pasteurella multocida* (accession code: HDR1349883.1) was synthesized by Twist Bioscience (South San Francisco, CA, USA). Each gene was cloned into the pET28a vector (Novagen, Wisconsin, USA) using the NdeI and XhoI restriction sites. The constructs encoded full-length Aca1 (residues 1-79), Aca12 (residues 1-140), or Aca14 (residues 1-66), each fused to an N-terminal hexa-histidine tag followed by a thrombin cleavage site. The recombinant plasmids were transformed into *Escherichia coli* BL21(DE3) competent cells. For each construct, a single colony was inoculated into 5 mL lysogeny broth (LB) medium containing 50 μg/mL kanamycin and incubated overnight at 37°C. The culture was subsequently used to seed 1 L of fresh LB medium supplemented with the same antibiotic and grown at 37°C until the optical density at 600 nm (OD_600_) reached 0.7-0.8. Protein production was induced with 0.25 mM isopropyl β-D-1-thiogalactopyranoside (IPTG), followed by incubation at 20°C for 18 h with shaking at 220 rpm.

Cells were harvested by centrifugation at 5, 300 × g for 15 min at 20°C and resuspended in lysis buffer (20 mM Tris-HCl pH 8.0, 500 mM NaCl, 1 mM phenylmethylsulfonyl fluoride). Cell disruption was performed by ultrasonication on ice, and the lysate was clarified by centrifugation at 38, 920 × g for 30 min at 4°C. The supernatant was incubated with Ni-nitrilotriacetic acid (Ni-NTA) affinity resins for 2 h at 4°C and loaded onto a gravity-flow column (Bio-Rad, Hercules, CA, USA). The resin was washed extensively with 50 mL lysis buffer to remove non-specifically bound impurities.

For removal of the N-terminal His-tag, the Ni-NTA resin-bound His-tagged protein was incubated with 80 U thrombin in cleavage reaction buffer (20 mM Tris-HCl pH 8.0, 100 mM NaCl, 0.3 mM CaCl_2_) for 4 h at 4°C. The untagged protein was collected from the flow-through fraction and further purified by size-exclusion chromatography (SEC) on a Superdex 200 Increase 10/300 GL column (GE Healthcare, Waukesha, WI, USA), pre-equilibrated with the corresponding buffer, 20 mM Tris-HCl (pH 8.0) and 500 mM NaCl for Aca1 and Aca14, or 20 mM Tris-HCl (pH 8.0) and 150 mM NaCl for Aca12. The elution profile was monitored, and fractions corresponding to the main peak were pooled and analysed by sodium dodecyl sulfate-polyacrylamide gel electrophoresis (SDS-PAGE) to confirm purity.

### RNA Electrophoretic mobility shift assays (EMSAs)

RNA EMSAs were performed based on a previously published protocol(11). Reactions consisted of purified Aca1 at the concentrations indicated in the figures, or no protein as a negative control, and 2 nM RNA in a buffer containing 0.1 mM TCEP, 25 mM HEPES-NaOH (pH 7.5) and 300 mM NaCl. RNA probes with IRDye® 800 label were obtained from Integrated DNA Technologies (IDT) and are listed in **Supplementary Table S6**. Reactions were incubated for 15 min in the dark, followed by addition of 2.5 µL loading dye (0.5× TBE (45 mM Tris (pH 8.3) 45 mM boric acid, 1 mM EDTA), 34% glycerol (v/v), 0.2% bromophenol blue (w/v)) and loading onto 8% polyacrylamide gels (19:1 acrylamide/bis acrylamide (Bio-Rad), 0.5× TBE, 2.5% (v/v) glycerol, 0.6 mg ml^−1^ ammonium persulfate, 0.05% (v/v) tetramethylenediamine)) that had been pre-run for 30 min at 4°C. Gel electrophoresis was performed at 150 V and 4°C in the dark for ∼1.5 h. RNA was imaged at 800 nm using the LI-COR Odyssey Fc imaging system and Image Studio software. Band intensities were quantified using GelAnalyzer 26.1 (available at www.gelanalyzer.com). Bands were initially detected using default parameters and manually adjusted to ensure consistent integration of free and shifted RNA species across lanes. The baseline was detected using the Rolling Ball method with a peak width tolerance of 50%. Intensities of all shifted RNA species were summed and expressed as a fraction of the total RNA signal. GraphPad Prism 11.0.2 was used to plot the fraction of bound RNA against protein concentration and a hyperbolic curve was fitted by nonlinear regression to estimate the apparent dissociation constant K_D_.

### DNA EMSAs

DNA EMSAs were performed using purified Aca12 and Aca14 proteins and FAM-labeled DNA probes corresponding to the predicted IR sequences identified in their respective promoter regions. For binding assays, 10 nM of the corresponding FAM-labeled DNA probe was incubated with increasing concentrations of Aca12 (50, 100, 250, 500, 1, 000, and 2, 000 nM) or Aca14 (10, 25, 50, 100, 250, 500, and 1, 000 nM) for 1 h at 25°C. For specificity analyses of Aca14, excess unlabeled specific or nonspecific competitor DNA (50 μM) was preincubated with 1, 000 nM Aca14 for 30 min at 25°C prior to the addition of 10 nM FAM-labeled DNA probe, followed by incubation for an additional 1 h at 25°C. Reaction mixtures were resolved on a 10% native PAGE gel (0.5× TBE) at 100 V for 1 h, and fluorescent signals were detected using an ImageQuant 800 imaging system (Cytiva, Marlborough, MA, USA).

### Reporter assays

Reporter assays were performed in *E. coli* DH5ɑ using a two-plasmid setup: one plasmid for arabinose-inducible expression of *aca14* from *Pasteurella multocida* strain P2095 (accession CP097618) and one plasmid with a transcriptional fusion of an *aca14*-associated promoter, or variant thereof, to *eyfp*. The expression plasmid was generated by inserting the *aca14* gene with an artificial ribosome binding site (synthesised as a gBlock from Integrated DNA Technologies) into pBAD30(42) between the EcoRI and HindIII sites. The *eyfp* reporter plasmids were generated by inserting the promoter sequences corresponding to the *aca14* or the downstream operon, including mutated variants, together with an unrelated 5′ UTR, into pPF1439(43) between the SpeI and NsiI sites. The constructed plasmids are listed in **Supplementary Table S4** and the exact sequences cloned in **Supplementary Table S5**. *E. coli* DH5α cells co-transformed with expression and reporter plasmids, or the corresponding empty-vector controls, were grown at 30°C in LB medium supplemented with ampicillin (50 μg/mL), chloramphenicol (12.5 μg/mL), and 0.2% (w/v) arabinose. Cultures were diluted 1:200 in phosphate-buffered saline and examined using a Cytek Aurora spectral flow cytometer and SpectroFlo software. A total of 10, 000 events were recorded per sample. The main population of cells was first gated based on forward- and side-scatter areas. Fluorescence emission of eYFP was detected in the B3 channel. For samples that contained a single population of cells with background-level eYFP fluorescence, we reported the median eYFP fluorescence intensity of all cells within this gate. In samples with above-background eYFP fluorescence, the eYFP-positive cells were typically accompanied by a population of cells that had apparently lost eYFP fluorescence. In these instances, we further gated on the eYFP-positive cells and reported the median eYFP fluorescence intensity of this population.

## Results

### Bioinformatic discovery of nucleotide motifs adjacent to *acr–aca* operons

We previously investigated the anti-CRISPR-associated protein Aca2 from the *acrIF8–aca2* operon of *Pectobacterium carotovorum* phage ZF40. Aca2 uses its HTH domain to autorepress transcription of its operon by binding an inverted repeat overlapping with the promoter, but also to inhibit AcrIF8 translation from its mRNA by binding conserved hairpins in the 5′ UTR(11, 16, 17). To explore whether other Aca proteins are associated with potential RNA-binding motifs, we developed a pipeline to identify conserved nucleotide motifs upstream (100 nt) of *acr–aca* operons (Methods) (**Figure 1A**). We performed a proof-of-concept analysis by analysing the RNA and DNA motifs of *acr–aca2* operons. In previous work, we manually identified two distinct RNA motifs, named after the bacterial phyla in which they predominantly occurred—the actino- and proteo-motifs(11). Using our computational pipeline, we detected 109 matches to these two motifs located upstream of *acr–aca2* operons across 98 distinct loci. Among these, 78 operons contained a high-confidence transcription start site prediction positioned downstream of the DNA motif. Both the proteo- and actino-motifs were represented within this high-confidence subset, demonstrating that the new pipeline reliably reproduced prior results in a high-throughput and systematic manner, thereby reducing manual curation.

**Figure 1:**
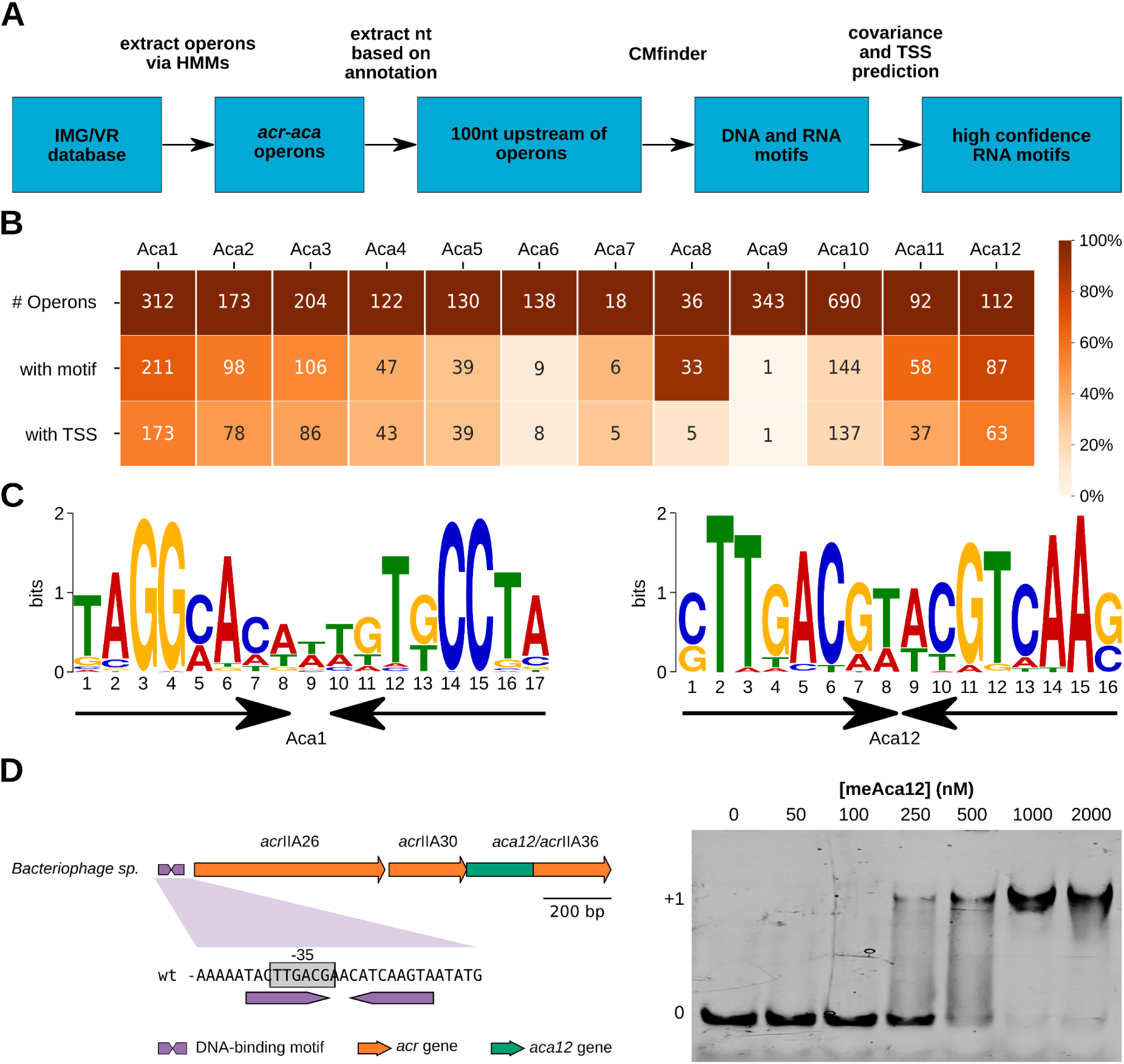
DNA motifs upstream of *acr–aca* operons. **A)** Summary of the discovery pipeline for potential Aca binding motifs. **B)** Number of analysed *acr–aca* operons and the number of operons with identified DNA motifs and predicted transcription start sites (TSS). Match to the DNA motif after one CM-based search with the E-value cutoff of 0.001. Colour code represents the percentage of operons (first row equals 100%). **C)** Predicted DNA binding sites of Aca1 (as previously described(10, 16, 17, 45)) and a novel predicted DNA binding site of Aca12. The black arrows indicate the inverted DNA repeats. The DNA motifs were visualised with the MEME suite(24). **D)** Overview of a *Bacteriophage sp.* operon encoding Aca12, AcrIIA26, and AcrIIA30. The predicted inverted repeat within the promoter is indicated. Nucleotide identifier: BK039523.1:12550..13900 (left). *In vitro* DNA binding by Aca12 of the predicted inverted repeat detected by EMSA. DNA was used at a concentration of 10 nM together with increasing concentrations of Aca12 as indicated. The dominant unshifted and shifted (protein-bound) DNA bands are denoted “0” and “+1”, respectively (right).

Following this validation, we applied the pipeline to operons (n=4673) of all other experimentally characterised Aca proteins with HTH domains (Aca1-Aca12). Our initial goal was to identify all known DNA motifs of Aca1 through Aca11 and to predict novel motifs for Aca12, for which no DNA binding site had previously been validated. The pipeline successfully recovered DNA motifs for 10 of the 12 Aca families, including a newly discovered inverted repeat associated with operons encoding Aca12 (**Figure 1B**). For Aca1–Aca11, the predicted motifs closely matched those established experimentally, supporting the accuracy and robustness of the pipeline(17, 19) (**Supplementary Figure S2**). To validate the newly predicted Aca12 motif (**Figure 1C**), we performed electrophoretic mobility shift assays (EMSAs) using the candidate inverted repeat. Aca12, which was previously described as AcrIIA36(44), produced a clear mobility shift, confirming binding to the predicted motif (**Figure 1D**). Together, these results validate the pipeline and show that it can reliably identify Aca-binding motifs upstream of *aca* operons across most Aca families.

### 5′ UTRs of *acr–aca1* operons contain RNA motifs for Aca1 binding

To identify conserved RNA motifs in the 5′ UTRs of *acr–aca* operons beyond those encoding Aca2, we focused on the region between the DNA motif and the start codon. We first investigated Aca1, since it is one of the Aca proteins with the most known homologs, shares operons with many different *acr* genes, and is found in phages of several different bacterial species (**Figure 2A**). In addition, its role in autoregulation via DNA binding has been thoroughly studied, and its high-resolution structure in complex with DNA has been solved(10, 45).

**Figure 2:**
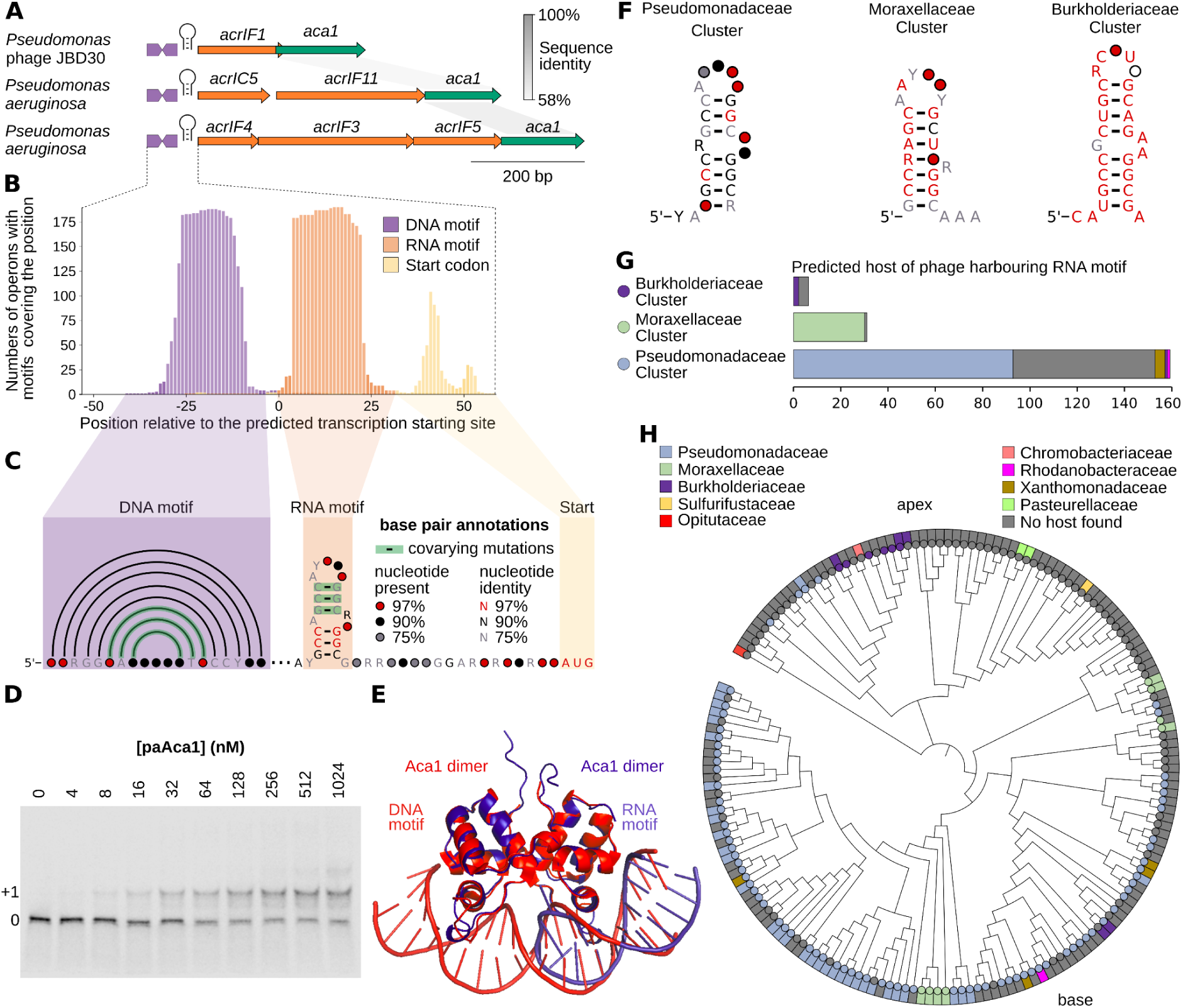
*acr–aca1* operons contain conserved RNA motifs in their 5′ UTR that are bound by Aca1. **A)** Three example operons encoding Aca1 (green) and Acr proteins (orange) with the respective DNA and RNA motifs (plum). Nucleotide identifiers: NC_020198.1:18750..193500, NZ_RABZ01000108.1:1822..2929, and NZ_JTMW01000030.1:17642..19417. **B)** Position plot from the 5′ UTRs of *aca1* operons indicating the position of each RNA motif (orange), DNA motif (blue), and start codon (yellow) relative to the predicted transcription start site. **C)** Consensus sequence and RNA secondary structure of the conserved DNA binding site (blue) and the predicted RNA motif (orange) from an alignment of 207 *acr–aca1* operons. Nucleotide colour indicates conservation, and the colour circles indicate how frequently any nucleotide is present at this position (if none of the four nucleotides occur more frequently than 75%). Black half-circles indicate complementary bases in the DNA motif. Green-shaded base pairs co-vary significantly. The three small squares denote the boundary between sequences that operate at the DNA or RNA levels. **D)** *In vitro* RNA binding by Aca1 detected by EMSA. RNA was used at a concentration of 2 nM together with increasing concentrations of Aca1 as indicated. The dominant unshifted and shifted (protein-bound) RNA bands are denoted “0” and “+1”, respectively. For uncropped triplicate gels, K_D_ plot and specificity controls, see **Supplementary Figure S3**. **E)** Predicted AF3 protein structure for the Aca1 dimer binding RNA (blue, right side, AF3 confidence scores can be seen in **Supplementary Figure S4**) in comparison to the crystal structure of Aca1 binding DNA (red, long double helix, PDB:7VJM). **F)** Consensus RNA secondary structures of the three RNA motif clusters. For base annotation see panel **C)**. **G)** The predicted hosts of the phages that harbour the *aca1* operon of each RNA cluster, termed the Burkholderiaceae cluster (top, violet), the Moraxellaceae cluster (middle, green), and the Pseudomonadaceae cluster (bottom, blue). Predicted host family assignments were taken from IMG/VR metadata. These clusters cover 195 out of 207 phages harbouring the RNA motif. **H)** Unrooted phylogenetic tree of 171 Aca1 proteins, with distances of nodes from the depicted root in arbitrary units that do not represent evolutionary time or strength of relatedness. Coloured circles on the outside indicate whether the RNA motif is present in the associated operon and to which cluster it belongs (colouring based on **G**). Coloured rectangles show the predicted host of the Aca1-encoding phage. Bootstrap values and branch length are shown in **Supplementary Figure S5**.

The architecture of the upstream region of *acr-aca1* operons is highly conserved, with the DNA-binding motif in the promoter region of the operon, and the predicted transcription start site around 10 nt downstream. In 86 out of 211 (∼40%) *acr-aca1* operons, an additional DNA motif, described previously, was identified upstream of the one overlapping the promoter(10). In addition, we identified one RNA motif upstream of 207 different *acr-aca1* operons. This motif is an inverted repeat (IR) of around 20 nt, lies directly downstream of the transcription start site (**Figure 2B, C**) and ends approximately 20 nt upstream of the *acr* start codon. The RNA hairpin predicted to be formed by this IR has a lower stem consisting of three highly conserved base pairs, which are separated by up to two unpaired bases from the upper stem. The upper stem has three base pairs, which show statistically significant covariation (**Figure 2C**).

To test if Aca1 directly interacts with the 5′ UTR containing the predicted RNA motif, we purified Aca1 from the *acrIF1–aca1* operon of *Pseudomonas aeruginosa* phage JBD30 and incubated it with a fluorescently labelled RNA probe containing the first 60 nt of the *acrIF1–aca1* mRNA. EMSAs with increasing Aca1 concentrations demonstrated binding to the *acrIF1–aca1* 5′ UTR RNA as a more slowly migrating product (**Figure 2D, Supplementary Figure S3A**), with an apparent dissociation constant (K_D_) of ∼61 nM (**Supplementary Figure S3B**), which is twice as high as the K_D_ of Aca2 binding its RNA motif. No binding was detected to a range of non-cognate RNA probes of equal length and with similar potential for stemloop formation, which suggests a specific interaction of Aca1 and its RNA motif (**Supplementary Figure S3C**). We followed the EMSA with structural predictions of the Aca1 dimer binding its respective RNA targets using AlphaFold 3. This prediction is in line with the experimental results and potentially suggests that the stem-loop formed by the bases of the RNA motif forms a double helix and could be bound by the HTH domain in a manner similar to its binding to DNA (**Figure 2E, Supplementary Figure S4**), reminiscent of our earlier finding with Aca2(11). These results demonstrate that the RNA motif identified in the *acr–aca1* 5′ UTR can be bound by Aca1.

### Aca1 is associated with host-specific RNA motifs

The interaction of Aca1 with its RNA motif in the 5′ UTR could result in co-evolution between the amino acid sequence of Aca1 and the RNA motif sequence. Similarly, our previous work detected these relationships with Aca2(11). To probe host-specific adaptation of Aca1–RNA pairs, we first clustered RNA motifs by sequence similarity and mapped these groups to the predicted hosts of these phages. Three clusters emerged: a large cluster (158 motifs) with diverse hosts primarily from the family Pseudomonadaceae, a mid-sized cluster (31 motifs) exclusive to Moraxellaceae, and a small cluster (6 motifs) specific to Burkholderiaceae. This suggests a potential correlation between the phage host and the RNA motif (**Figure 2F–G**).

Next, we analysed Aca1 phylogeny and annotated the respective RNA motif clusters on the resulting tree (**Figure 2H**). The tree revealed clear concordance: Aca1 proteins from Pseudomonadaceae-associated phages formed a clade at the base of the tree, while Aca1-encoding Burkholderiaceae-targeting phages clustered distantly at the apex. This could indicate a non-random association between Aca1 sequence relatedness and upstream RNA-motif identity. To quantify this pattern, we treated identity in one of three RNA clusters as a discrete trait on the fixed Aca1 tree and calculated the Fitch parsimony score, for which lower values indicate fewer state changes across the tree. The observed parsimony score was significantly lower than expected from randomized tip-label permutations, consistent with phylogenetic clustering of RNA motif classes on the Aca1 tree (**Supplementary Table S1**). The association between Aca1 phylogeny and RNA cluster identity remained significant in the unclustered dataset and after dereplication at 95% and 90% amino-acid identity. Together, these analyses support a stable lineage-associated relationship between Aca1 protein diversification and the identity of the linked 5′ UTR RNA motif. In contrast, the Moraxellaceae cluster is split into two groups. This divergence may reflect increased phylogenetic sensitivity to minor amino acid changes in the small Aca1 protein (<100 residues), where small changes in the sequence can disproportionately influence branching patterns. This phylogenetic analysis is consistent with the hypothesis that the experimentally validated protein–RNA interaction is an evolutionarily conserved aspect of its function.

### Operons containing *aca10* and *aca12* have predicted RNA motifs

To examine if Aca proteins beyond Aca1 and Aca2 recognise RNA motifs, we performed a bioinformatic search across the operons containing the remaining *aca* genes (i.e., *aca3* to *aca12*). Potential motifs were identified in the *acr–aca10* and *acr–aca12* operons (**Supplementary Figure S6**). Although other *aca* operons contained predicted RNA motifs in their predicted 5’ UTR, their covariation signal was insufficient to assign these as biologically relevant RNA motifs, which is likely due to insufficient diversity of available sequences. We examined Aca10 and Aca12 for phylogenetic evidence of co-evolution between the protein and the RNA motif; however, no correlation was detected. For Aca12, this absence of correlation likely stems from its exclusive presence in phages infecting diverse *Streptococcaceae* hosts, precluding lineage-specific motif adaptation.

Both the Aca10- and Aca12-associated RNA motifs show statistically significant covariation in their predicted RNA hairpins and, for Aca10, in the DNA motif. The upstream regions of *aca10* and *aca12* operons with associated RNA motifs also have a similar architecture as the *aca1* and *aca2* operons: the DNA motif overlaps with the predicted promoter region, the predicted transcription start site is 10 to 15 nt downstream of the DNA motif, and the RNA motif starts directly downstream of the transcription start site and is therefore predicted to be in the 5′ UTR. The position of the *acr* start codon of these operons is also highly conserved. To directly test for RNA binding, EMSAs similar to those used for the Aca1–RNA and Aca2–RNA interactions were performed with purified Aca10 and Aca12 proteins but have so far not shown binding to their predicted RNA motifs. Nevertheless, our bioinformatic and Aca1 results suggest that other Aca proteins could have additional functions in RNA binding – akin to the well-characterised Aca2 protein(11).

### Aca proteins are predicted to co-regulate multiple anti-defense operons

Phages commonly encode multiple Acr and other anti-defence proteins that likely require coordinated expression(46). We hypothesised that individual Aca proteins regulate multiple anti-defence genes and operons, with our computational identification of Aca DNA and RNA binding motifs providing an opportunity to test this hypothesis. To investigate multi-target regulation, we searched for bacterial and phage contigs encoding a single Aca protein family (e.g., Aca1) alongside multiple predicted binding sites for that regulator. Analysis of predicted Aca1 and Aca2 binding sites revealed instances where up to three predicted regulatory motifs clustered near operons in close genomic proximity. In each case, the Aca protein appeared to regulate gene clusters adjacent to the primary *aca*-containing operon. For example, in *Marinomonas spartinae*, three pairs of Aca2 DNA- and RNA-binding sites were identified upstream of three distinct neighboring operons, only one of which contained *aca2* (**Figure 3A**). Additionally, we identified RNA motifs positioned between adjacent genes (including known *acr* genes) within predicted individual operons, suggesting a mechanism for fine-tuning and tightly regulating translation of specific genes within anti-defence operons (**Figure 3A**).

**Figure 3:**
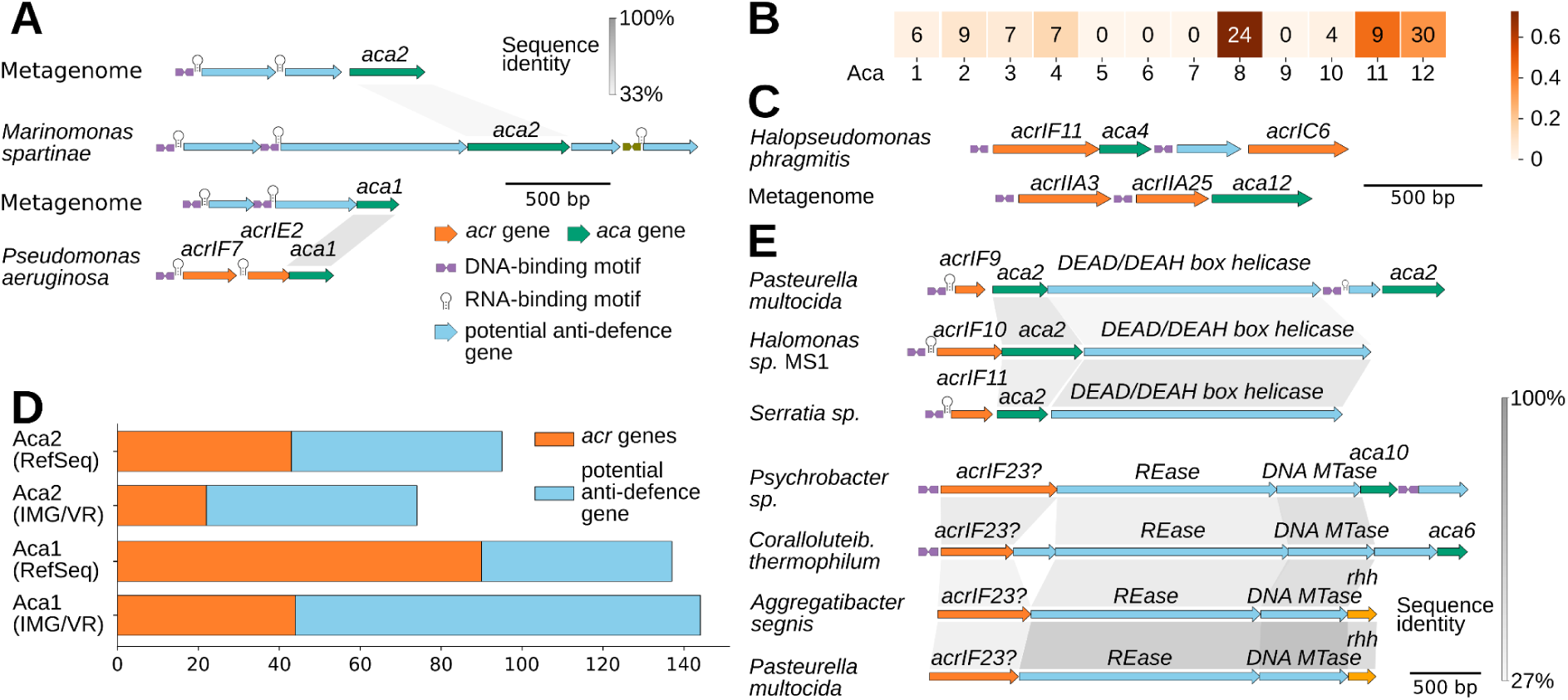
Aca proteins are predicted to regulate not only their operon but also other nearby anti-defence operons. **A)** Schematic of different Aca1 and Aca2 loci in (pro)phage regions. The DNA binding sites are indicated by the plum arrows, the RNA binding site is marked with the hairpin, *aca* genes in green, *acr* genes in orange, and other potential anti-defence genes in light blue. Nucleotide identifiers: Ga0326295_1000217: 12400..14100, NZ_FLOB01000002.1: 620600..623200, Ga0395800_0000105: 24800..26050, and NZ_SDVE01000001.1: 441450..442300. **B)** Number of Aca proteins predicted to regulate at least one other operon other than their own. Proteins were counted if a DNA motif was present upstream of a different operon. Heatmap colouring is based on the percentage of Aca proteins regulating more than one operon. **C)** Synteny plot for two example Aca (Aca4 and Aca12) proteins regulating more than one operon and colouring according to Figure 3A. Nucleotide identifiers: CP020100.1: c2620200..2618650 and Ga0099518_101633: 2600..3900. **D)** Aca1 and Aca2 regulate many genes without known function. Number of genes with regulatory DNA and RNA motifs of Aca1 and Aca2 upstream in the RefSeq and IMG/VR database. The number of orange (left) genes is annotated as *acr* genes, and blue (right) genes could be potential anti-defence genes. **E)** Three different *acr-aca2* operons with a predicted DEAD/DEAH box helicase downstream of *aca2* in three different bacterial hosts. The helicase is predicted to be in the same operon and regulated by the Aca2 protein (upper part). Restriction-Modification systems encoding restriction endonucleases (REases) and methyltransferases (MTases) can be embedded in *acr-aca* operons, and the typical HTH-domain Aca can be substituted by the RHH-domain protein (lower part). Distant homologues of AcrIF23 are marked with a question mark. Colouring according to Figure 4A. Nucleotide identifiers: NZ_JAMJUV010000016.1: 2400..6087, NZ_CP098926.1: c3808500..3811700, NZ_CP109901.1: 1083800..1086800, NZ_JBEOOX010000005.1: 151401..155201, NZ_JBHSGG010000036.1: c331..4231, NZ_NRCV01000004.1: 159307..165516, and NZ_AP025519.1: 886500..890800.

To ascertain whether the predicted regulation of neighbouring operons is a common feature of Aca proteins, we conducted a search of our dataset of predicted DNA-binding motifs (E-value cutoff 0.001). With the exception of Aca5 and Aca7, we identified examples with the potential to regulate nearby operons in all Aca families (**Figure 3B, 3C, and Supplementary Figure 7**). Interestingly, Aca11 and Aca13 were recently shown to regulate not only their own *acr–aca* operons but also adjacent downstream operons(19). Together with our results, this suggests that most, if not all, Aca proteins may extend their regulatory influence to neighbouring operons.

### Aca-regulated operons encode putative novel anti-defence proteins

We hypothesised that uncharacterized genes predicted to undergo Aca regulation also encode anti-defence proteins. To test this, we used DNA and RNA motifs for Aca1 and Aca2 to assess the proportion of annotated *acr* genes relative to putative anti-defence candidates located downstream of these regulatory elements. Integrating data from RefSeq and IMG/VR databases, we identified 450 genes under predicted Aca1 and Aca2 regulation, with fewer than half having assigned functions (**Figure 3D**). Similar distributions were observed for genes directly downstream of other Aca regulatory sequences (**Supplementary Figure S8**). To examine whether some Aca-regulated genes have broader anti-defence capabilities, the anti-prokaryotic immune system database (dbAPIS)(36), the conserved domain database (CDD)(35), and the DefenceFinder with the anti-defence add-on(37) were employed for the analysis of the proteins in operons that are predicted to be regulated by Aca proteins. Among the nearly 1, 900 genes predicted to be regulated by Aca1-12, there were over 420 *acr* genes and 666 hits to the CDD. As this included the *acr–aca* operons themselves, more than 500 proteins had a match to either Acr or HTH domains. However, multiple other domains were present in comparably high numbers as Acr proteins, suggesting that Aca proteins regulate diverse genes potentially involved in immune inhibition and mobile genetic element conflicts. In addition, we identified multiple protein clusters lacking annotated domains or assigned functions, which may represent anti-defence candidates for future investigation (**Supplementary Figure S9**).

The most frequently occurring conserved domain identified in *acr–aca2* operons after DUF1870 (the HTH domain in Aca2) and AcrIF9 was the COG4889 and HA superfamily, both as part of the same protein. COG4889 is a predicted helicase domain, while HA is a helicase-associated domain and can be found near Retrons(47). The proteins containing these domains were annotated as DEAD/DEAH box helicases by CDD (**Figure 3E**). These proteins can be involved in eukaryotic and prokaryotic immune systems(48, 49).

While searching for anti-defence candidates in *acr–aca* operons, we found several examples of type II restriction–modification (RM) systems embedded between a distant homolog of *acrIF23* and the *aca* genes (**Figure 3E**). The distant *acrIF23* homolog was frequently associated with helicases and nucleases (**Supplementary Figure S10**). In addition, we detected an apparently Aca-regulated homolog of the phage Mu Mom protein, which is involved in DNA hypermodification(50), and several instances of phosphoadenosine phosphosulphate (PAPS) reductases, which are predicted to be homologs of DndC and SspD (**Supplementary Figure S10**). Notably, DndC and SspD are key components of the DNA phosphorothioate (PT) modification system DndABCD and SspBCDE(51–53), and they also exhibit PAPS reductase activity. Since both Mom and PAPS reductases could participate in DNA modification, they may represent host-derived components repurposed by phages to evade RM or PT systems(51, 52, 54, 55). Consistent with this idea, PAPS reductase proteins cluster in variable phage genomic regions, which are often hotspots for anti-defence genes(44). In addition, previous work has shown that a phage-encoded PAPS reductase can promote evasion from the host Dnd system(56).

We also detected other conserved domains. For example, four genes encoding LPD29 (Large polyvalent protein associated) domains were present in *aca4* operons. Polyvalent proteins are proposed to be part of the intergenomic conflict between MGEs and are found in the leading regions of phages and plasmids(57). LPD proteins are bioinformatically predicted to inhibit bacterial defence systems, e.g. RM, suggesting that LPD29 could also have an anti-defence function(57). In summary, our data suggest that Aca proteins regulate not just *acr* genes but also different anti-defences or other factors that could provide a competitive advantage against the host or superinfecting phages. Although most Aca-associated genes remain unannotated, our functional predictions suggest that a substantial fraction are involved in counter-defence mechanisms. This association and the identified protein clusters provide a framework for prioritising these genes in future anti-defence functional assays.

### A novel ribbon-helix-helix Aca protein regulates multiple operons

We searched within the non-redundant protein database for proteins similar to the regulated RM systems just described to identify additional *acr–rm–aca* operons. This search revealed an operon in which the *aca* gene is replaced by a gene encoding a ribbon-helix-helix (RHH) domain protein (**Figure 3E**). As RHH domains are frequently found in transcription factors that autoregulate or control other operons(58), this gene may encode a novel candidate Aca protein. We searched the PSI-BLAST database for homologs of this RHH protein and found several representatives encoded within *acr* operons (**Figure 4A**), most frequently associated with AcrIF23, AcrIF6, AcrVA3 in operons lacking known *aca* genes. Since RHH transcription factors typically recognise inverted DNA repeats(58), we applied our nucleotide motif discovery pipeline to identify potential DNA-binding motifs for the RHH-domain proteins. This analysis revealed a highly conserved inverted repeat overlapping the predicted -10 promoter element (**Figure 4B and C**) but no conserved RNA motif with statistically significant covariation. These repeats were observed not only upstream of *acr–rhh* and *rm–rhh* operons but also in the upstream regions of neighbouring operons, mirroring the predicted regulatory versatility of Aca proteins. Moreover, we identified over 40 known *acr* genes in the operons predicted to be regulated by the RHH protein (**Supplementary Figure S11**). Interestingly, the predicted DNA-binding motif is also found upstream of several *csx20* homologs (**Figure 4A and Supplementary Figure S7**). Csx20 is a CRISPR-associated ring nuclease that cleaves cyclic oligoadenylates(59) and could be another example of a *cas* gene co-opted by phages for counter-defense(55), similar to the previously described AcrIII-1(60). Overall, predicted *acr–rhh* operons appear to be as common as *acr-aca12* operons.

**Figure 4:**
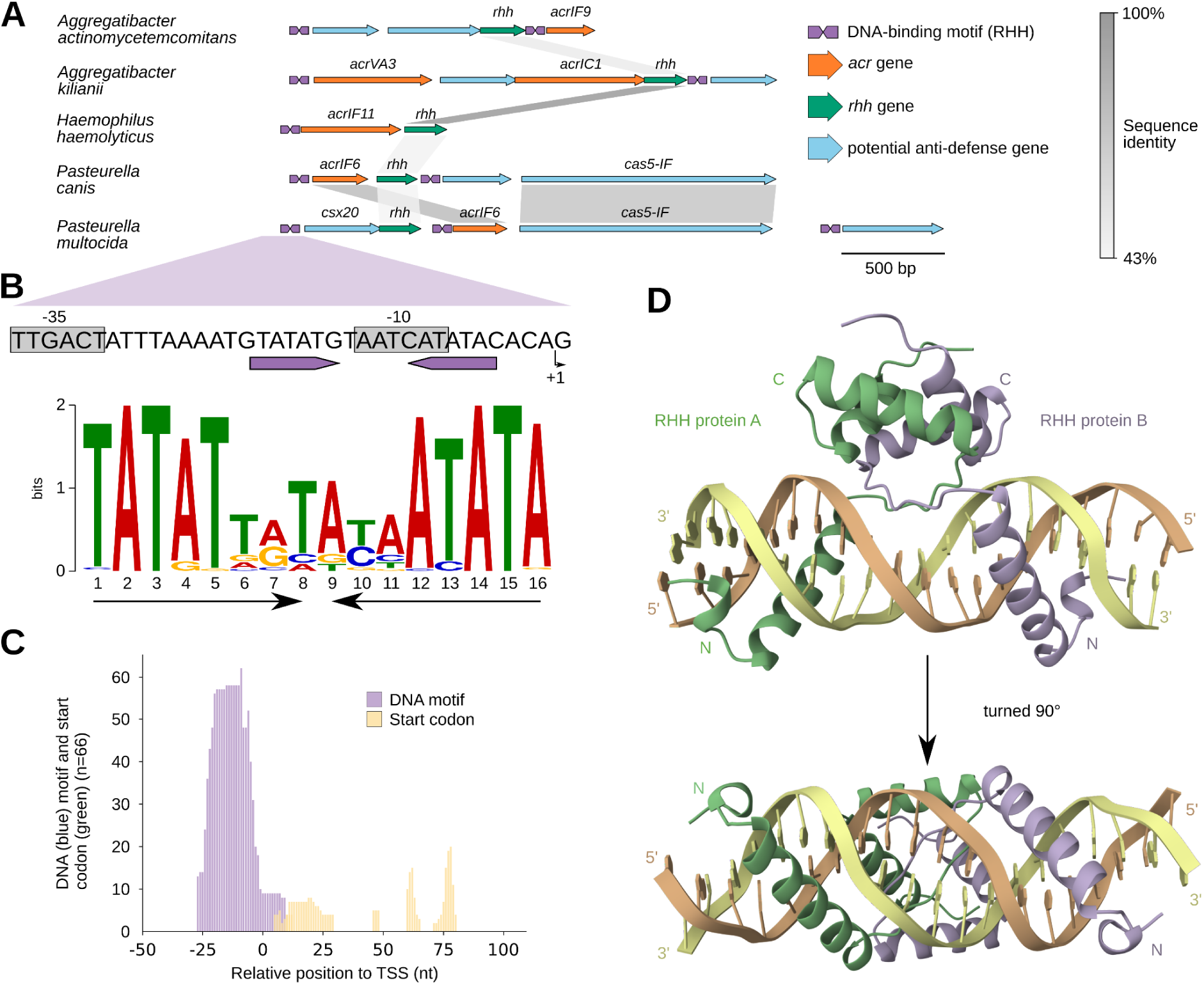
Anti-defence operons predicted to be regulated by an RHH-domain containing protein. **A)** Synteny plot of five different *acr* loci with an RHH-domain protein. Orange: *acr* genes, green: *rhh* gene, light blue: potential anti-defence gene, and plum arrows: predicted DNA motif of RHH protein. Nucleotide identifiers: NZ_AEJP02000019.1: 55000..58800, NZ_CAUQLS010000011.1: 29743..32200, NZ_QQIB01000001.1: 492600..493800, NZ_WUMP01000004.1: 88900..91300, and DAOHFP010000061.1: 0..3700. **B)** Promoter region with the DNA motif as predicted by the nucleotide motif discovery pipeline and visualised by MEME. Black arrows indicate inverted repeat. **C)** Relative position of the predicted DNA motif to the predicted transcription starting site of the *acr* operon. Blue: DNA motif and green: Start codon. **D)** Predicted structure of the RHH-domain protein binding the DNA motif by AF3 (Confidence scores can be seen in **Supplementary Figure S12**). The RHH-domain protein is predicted to form a dimer and insert itself in the major groove of the DNA.

Next, we tested whether an RHH protein could act as an Aca via interaction with these IRs. First, we used AlphaFold 3 to model the binding of an RHH Aca candidate, co-encoded in an operon with *csx20* in the pathogen *Pasteurella multocida* (**Figure 4A**), to its predicted AT-rich cognate IR DNA. AlphaFold 3 predicted a high-confidence model only for a homodimeric RHH complex bound to its associated inverted repeat (**Figure 4D, Supplementary Figure S12,** and **Table S2**), consistent with prior structural reports of RHH proteins(49). Notably, the model suggests that, in addition to the canonical RHH DNA-binding interface, a long helical extension inserts into the major groove of the DNA. When we purified this RHH protein, its elution volume during size-exclusion chromatography supported a dimeric state (**Figure S13**). We next used an EMSA to directly test binding of the purified protein to its cognate DNA motif. The RHH protein bound the predicted IR DNA in a concentration-dependent manner but showed no detectable binding to nonspecific competitor DNA (**Figure 5B, C**). To test if DNA binding results in autorepression of the *csx20–rhh* operon, we cloned the operon promoter in front of *eyfp* and performed flow cytometry to measure fluorescence in the presence and absence of RHH expression from a separate plasmid (**Figure 5D**). Expression of RHH repressed the reporter, but not when the IR sequence was mutated (**Figure 5E**). Based on our finding that this protein can repress its operon through specific interactions with binding sites in the promoter, we designate it Aca14. This family distinguishes itself from previously described anti-CRISPR-associated proteins by its use of an RHH rather than the canonical HTH domain.

**Figure 5:**
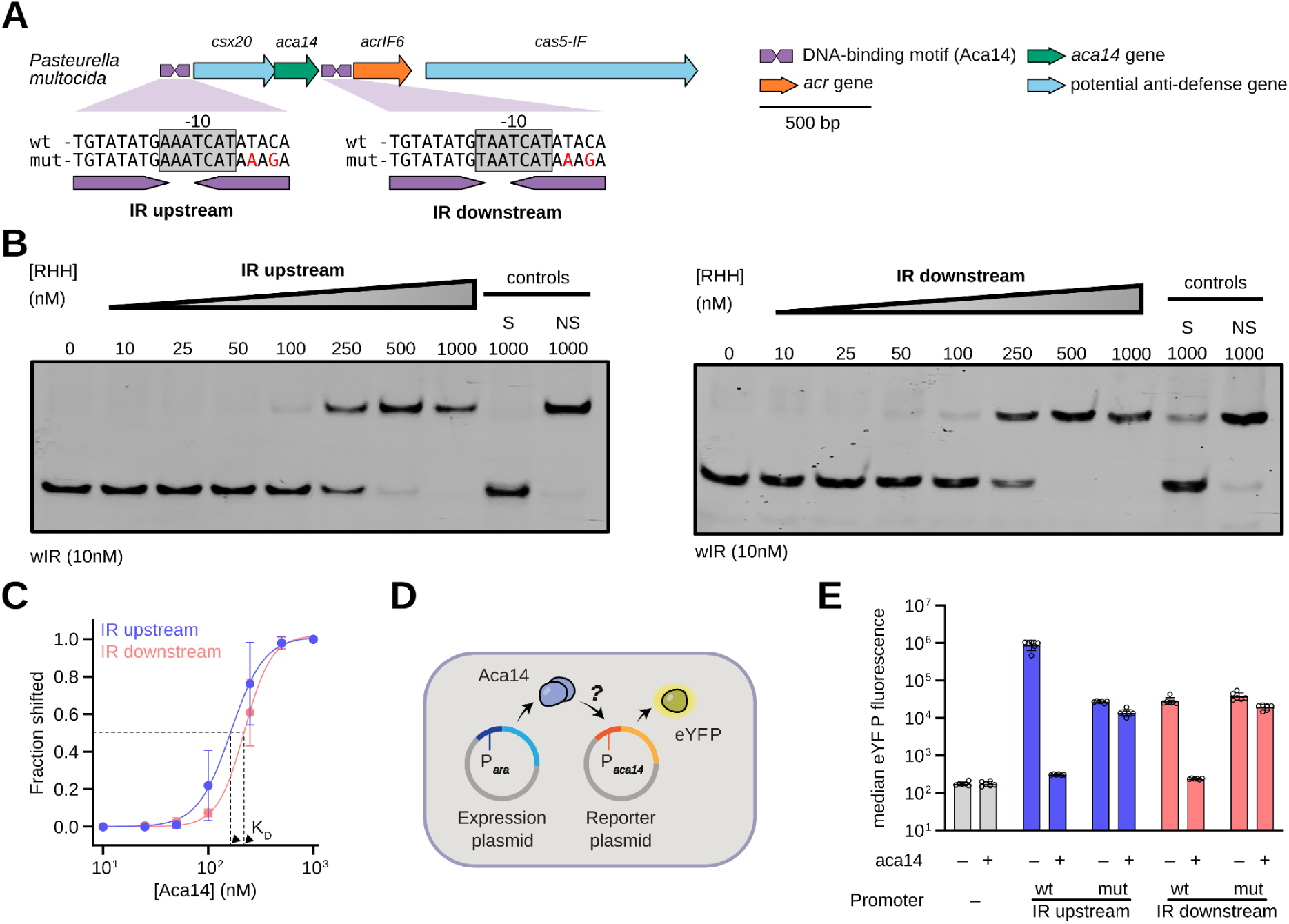
Experimental validation of the ribbon-helix-helix protein Aca14. **A)** Overview of a *Pasteurella multocida* operon encoding the predicted regulator Aca14 and a Csx20 homolog. An inverted repeat within the promoter is indicated. A similar site is found in the promoter of the downstream operon, which encodes homologs of AcrIF6 and Cas5-IF. ‘mut’ indicates mutated IR variants examined in the reporter assay in **E**. **B)** *In vitro* binding of Aca14 to the IRs (upstream and downstream of the *csx20–aca14* operon) with increasing Aca14 concentrations. Specific and non-specific controls are denoted S and NS, respectively. Representative examples from three independent experiments are shown. **C)** Dose-response curves based on the EMSAs in **B**, indicating the K_D_ for up- and downstream IRs (164 nM and 221 nM, respectively) with dashed lines and black triangles. **D)** Schematic of the reporter assay for testing Aca14 activity, involving an expression plasmid for arabinose-inducible Aca14 production and a reporter plasmid with *eyfp* fused to a predicted Aca14-controlled promoter. **E)** Results of the reporter assay testing reporter plasmids with the up- and downstream promoters predicted to be controlled by Aca14. Wild-type IR sequences and mutated variants are shown in panel **A**.

Like other Acas, the *aca14*-containing operon from *P. multocida* was located directly upstream of another putative anti-defence operon encoding AcrIF6 and a Cas5f homolog. The promoter of that operon overlapped with a predicted binding motif that was highly similar to that in the *aca14* operon promoter (**Figure 5A, Supplementary Figure S7**). This presented an opportunity to experimentally test our previous prediction that Aca proteins can regulate multiple operons (**Figure 3**). We adapted our EMSA and reporter assay to investigate whether Aca14 can also interact with the IR found beside the downstream operon to mediate repression. Indeed, DNA containing the binding motif from the downstream operon was specifically bound by purified Aca14 in an EMSA (**Figure 5B**), and both interactions had similar dissociation constants (**Figure 5C**). In addition, Aca14 repressed the downstream promoter using reporter assays and was dependent on an intact IR binding motif. These results provide strong evidence that Aca14 is a regulator not just of its own operon but also of the neighboring operon.

## Discussion

Here, we set out to explore the regulatory landscape of Aca proteins. In addition to their thoroughly investigated function as transcriptional regulators(10, 11, 17, 19), we show that roles in RNA-mediated regulation, as previously observed for Aca2(11), may also occur in other Aca families. Furthermore, our evidence suggests that the regulatory reach of Aca proteins frequently extends beyond their own operons to genes or operons encoding other defence and anti-defence components. One example is the newly discovered Aca14, which represents a family of anti-defence regulators with an RHH instead of the canonical HTH domain. Together, these findings portray Aca proteins as versatile tools for the coordinated and controlled production of diverse phage-encoded proteins, highlighting their critical role in the arms race between phages and bacteria.

We previously demonstrated that the Aca2 dimer binds two conserved hairpins in the 5′ UTR of its operon’s mRNA, which leads to translational repression(11). Here, we showed that Aca1 also specifically binds a conserved RNA motif in its operon’s 5′ UTR. Unlike the Aca2 RNA motif, the Aca1 motif consists of only one stem-loop, and modelling suggests it interacts with one monomer from the Aca1 dimer. It is therefore possible that a single Aca1 dimer binds two RNA molecules. The exact effect of Aca1 binding to this motif is unknown as we observed no translational silencing in reporter assays (data not shown). The Aca1 RNA motif may also titrate Aca1 away from the promoter DNA motif to fine-tune gene expression, similar to the proposed role of a second DNA binding site upstream of the *acrIF1-aca1* promoter(10). Alternatively, binding could affect mRNA stability and turnover(61, 62). Together with previous findings showing that HTH-domain transcription factors can bind mRNA and modulate its turnover (63), these observations suggest that RNA binding is a shared feature of many HTH regulators.

In addition to their versatility regarding nucleic acid targets, our data indicate that Aca proteins can extend their regulatory scope beyond their own *acr–aca* operons to include other anti-defence and host-counteracting genes. Such coordinated regulation may be advantageous, as different Acrs and inhibitors of other host defences likely share similar expression requirements: strong and immediate induction following infection to suppress host immunity, followed by sustained moderation to minimise toxicity or unnecessary expression(10, 11, 64). Hence, regulation by a single Aca could synchronise the transcriptional response of functionally linked operons. This model is consistent with previous reports showing that Aca11 and Aca13 bind within the promoters of both their cognate and downstream operons(19). Our findings further suggest that Aca proteins control operons encoding diverse components such as DEAD/DEAH helicases, DNA modification enzymes, restriction enzymes, and LPD family proteins, thereby integrating regulation of multiple defence- and anti-defence-related functions. The phage-encoded DEAD/DEAH helicase identified here could, for example, interfere with host defence pathways by competing with host helicases for binding partners or cofactors(47–49). More broadly, our data raise the possibility that Aca-dependent control extends beyond DNA modification proteins to additional RM-associated components. Such co-regulation could provide a selective advantage by coupling immune evasion with suppression of competing mobile genetic elements, thereby reducing the risk of superinfection(46, 65–67). A similar regulatory logic has recently been proposed for archaeal viruses, in which early anti-defence and virus-encoded genes are predicted to be co-regulated by Aca8 binding motifs(46). Together, these observations imply that Aca proteins form a conserved regulatory network that coordinates the temporal expression of early phage genes involved in immune evasion across diverse bacterial and archaeal systems.

Our discovery of RHH domain proteins encoded in *acr* operons akin to *aca* genes, alongside predicted binding motifs, suggested that DNA-binding domains other than the HTH fold can regulate phage counter-defence. Indeed, we confirmed that a representative of this family of RHH regulators, which we coined Aca14, directly interacts with its predicted binding sites to repress the promoter of not just its own but also a neighboring operon. Interestingly, a recent study identified an RHH domain as part of the AcrIF25 protein, which was not essential for Acr activity and was replaced by an HTH domain in another AcrIF25 homolog(68). We previously speculated that this domain may represent a built-in regulatory module(69) similar to the HTH domains found in some dual-functional Acr proteins (70–72). Our finding of a role for RHH domain proteins in anti-defence regulation supports this idea. More generally, this work shows that a focus on HTH regulators may not be sufficient to uncover the whole diversity of the phage anti-defence landscape and that other regulatory domains can be co-opted for the same purpose. This raises the broader question of whether further DNA-binding domains commonly found in transcriptional regulators, such as zinc finger or helix-loop-helix(73, 74), may also function in *acr* operon regulation.

This study suggests that dual DNA- and RNA-binding by HTH-containing Aca proteins may be a conserved regulatory strategy across diverse anti-CRISPR systems. The identification of RHH proteins in canonical aca positions further indicates that phages have co-opted multiple structural scaffolds to control counter-defence operons. In addition, the many uncharacterized Aca-regulated genes point to a broader and largely unexplored repertoire of phage immune-evasion mechanisms. Together, these findings establish Aca and related proteins as adaptable regulators of phage counter-defence .

## Supplementary Data

Supplementary data can be found in the Supplementary Information file.

## Acknowledgments

The authors would like to thank the Center for Information Services and High-Performance Computing (ZIH) at TU Dresden for providing computer time. We gratefully acknowledge use of the Otago Micro and Nanoscale Imaging (OMNI) Flow Cytometry unit.

## Contributions

Conceptualisation: MF, NB, CMB, PCF and ZW; Methodology: MF and NB; Software: MF; Formal analysis: MF and NB; Investigation: MF and NB; Resources: CMB, SYL, HHP, CLB and PCF; Data Curation: MF and CMB; Writing - Original Draft: MF; Writing - Review & Editing: MF, NB, CLB, ZW and PCF; Visualisation: MF and NB; Supervision: PCF, HHP, CMB and ZW.

## Data availability

The data underlying this article are available in the GitHub repository: An expanded realm of anti-CRISPR-associated proteins and regulatory mechanisms, at https://github.com/MaxFeussner/An-expanded-realm-of-anti-CRISPR-associated-proteins-and-regulatory-mechanisms

