## Supplementary Figures and Tables for "An expanded realm of anti-CRISPR-associated proteins and regulatory mechanisms"

Supplementary Information

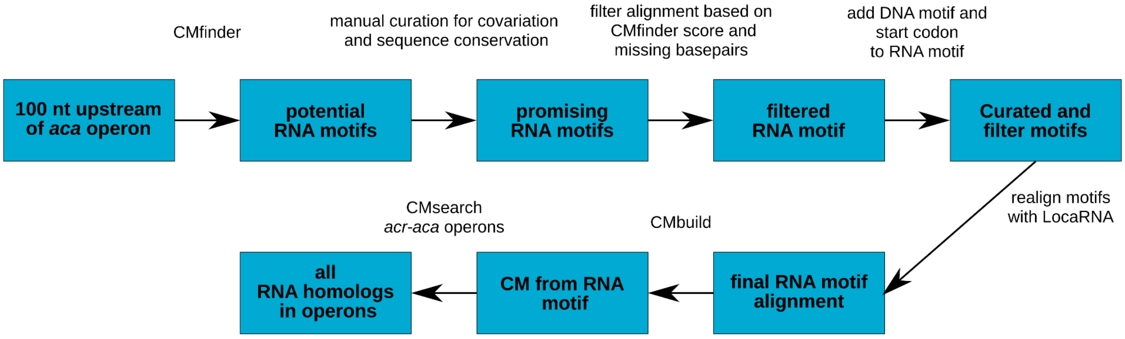


**Figure S1: Pipeline to identify conserved RNA and DNA motifs in front of *acr-aca* operons**. The pipeline uses the CMfinder software package[1] to identify novel RNA motifs in the 5′ UTR of *acr-aca* operons. Once a promising motif was identified, the Infernal software package was used to analyse the conservation of the motif within all *acr-aca* operons.

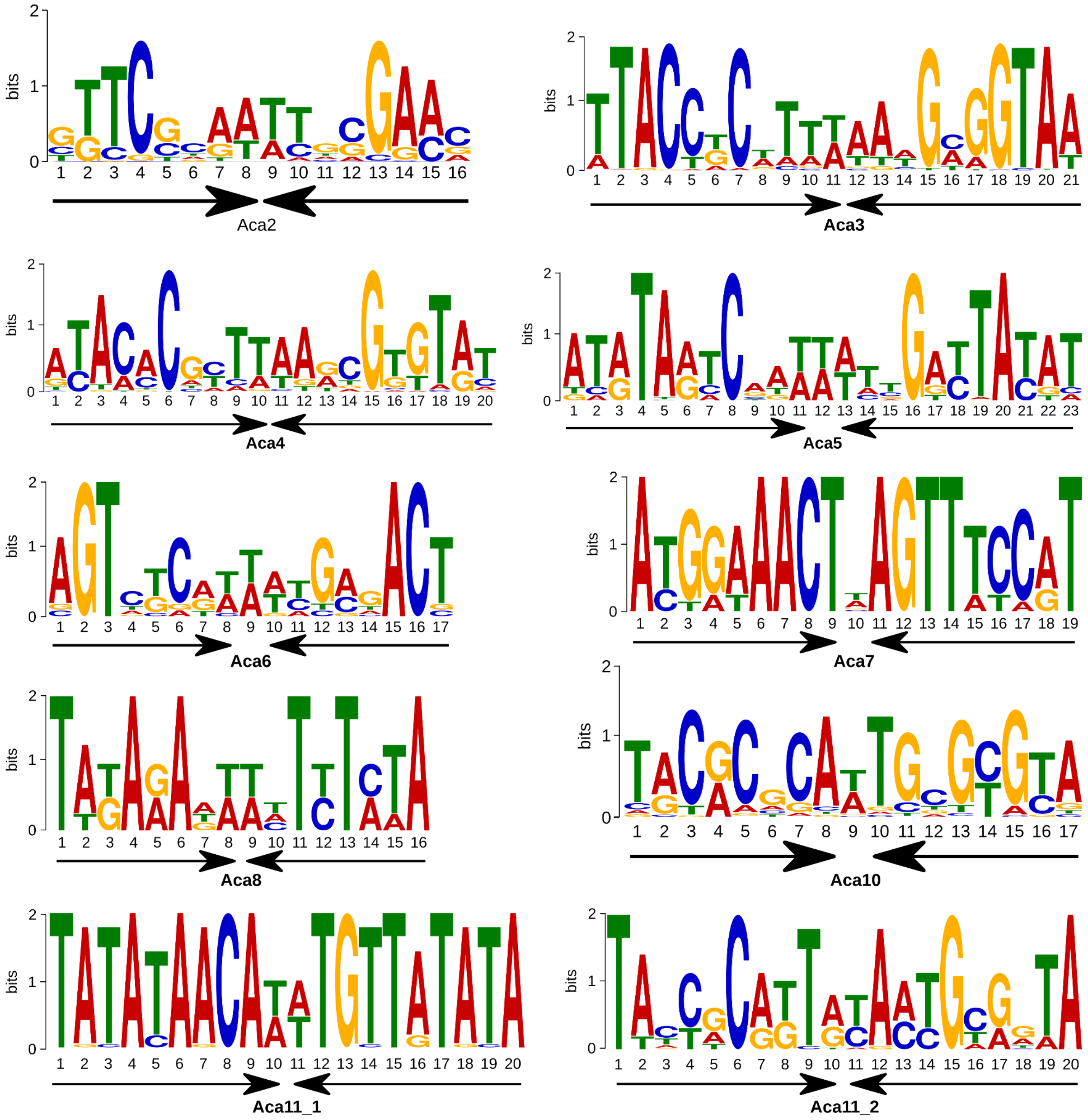


**Figure S2: Predicted DNA motifs of the Aca proteins**. The motifs were generated with the MEME suite[2] based on the DNA motifs with TSS downstream (**Figure 1B**). Black arrows indicate inverted repeats. Although MEME suggested two distinct DNA motifs for Aca11, our pipeline classified them within the same motif group. Motifs for Aca1, Aca2, Aca10 and Aca12 were described or referenced in the text, and are not shown here.

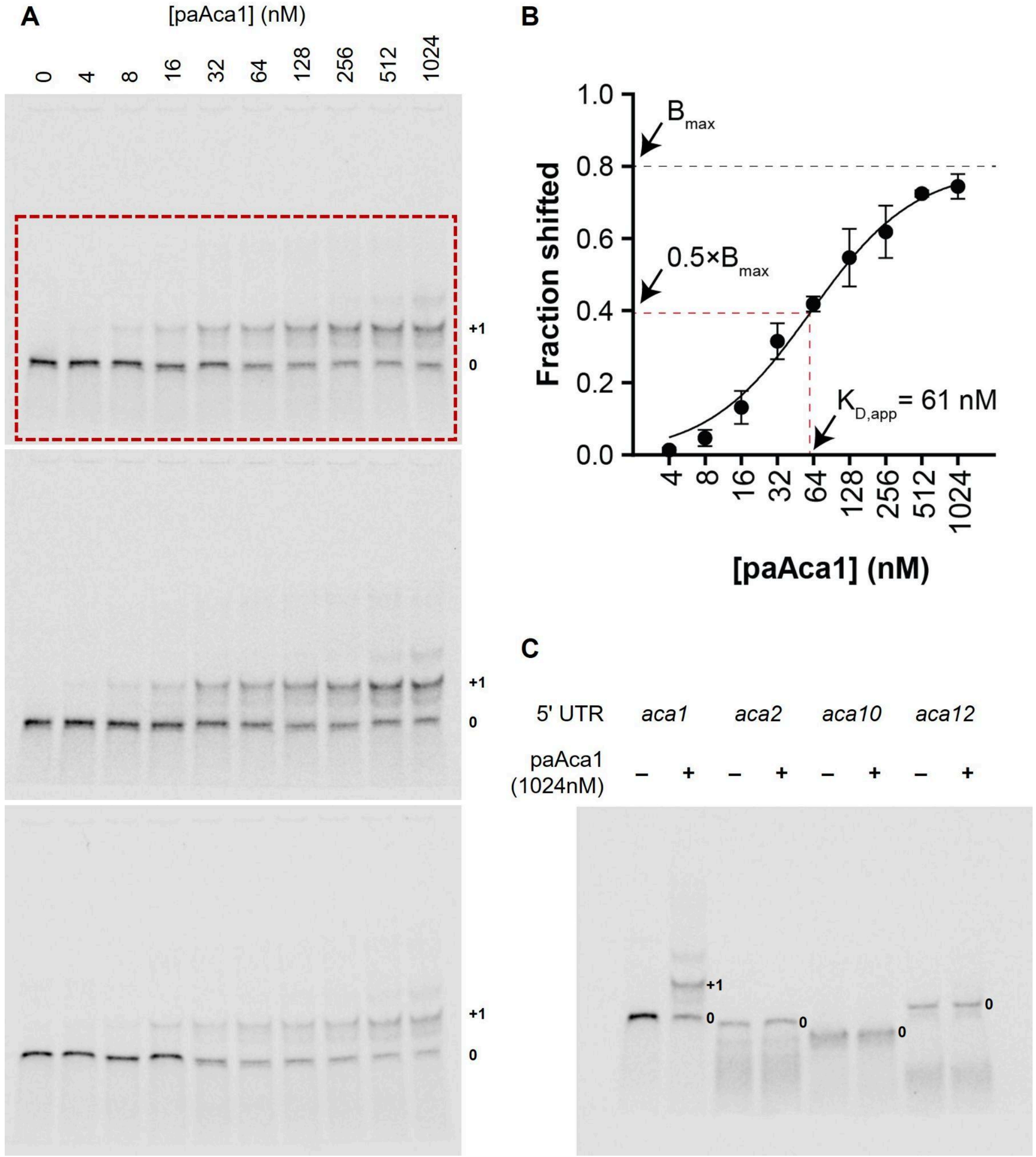


**Figure S3: Quantification and specificity of Aca1–RNA gel shifts. (A)** Triplicate uncropped EMSA gels of 2nM fluorescent RNA with the first 60nt of the *acrIF1–aca1* 5′ UTR (PF8279) with the indicated concentrations of purified Aca1 from *Pseudomonas aeruginosa* phage JBD30 (paAca1). The red dashed box indicates the gel portion shown in **Figure 2D**. The dominant unshifted and shifted (protein-bound) RNA bands are denoted “0” and “+1”, respectively. **(B)** Quantified fractions of shifted RNA for the three replicates as a function of Aca1 concentration. $B_{max}$ indicates the maximum fraction bound and $K_{D,app}$ the apparent dissociation constant, i.e. the Aca1 concentration at $0.5{\times}B_{max}$. All concentrations refer to monomeric Aca1. **(C)** EMSA of purified paAca1 after incubation with different RNA probes as specificity controls. Aca1 was used at 1,024nM, the maximum concentration used in the EMSAs in **(A)**. Specificity controls are 5′ UTRs from an *aca2*-associated operon from

*Pectobacterium carotovorum* phage ZF40 (PF4483); an *aca10*-associated operon from *Pseudomonas citronellolis* (PF9678); and a metagenomic *aca12*-associated operon (PF9679).

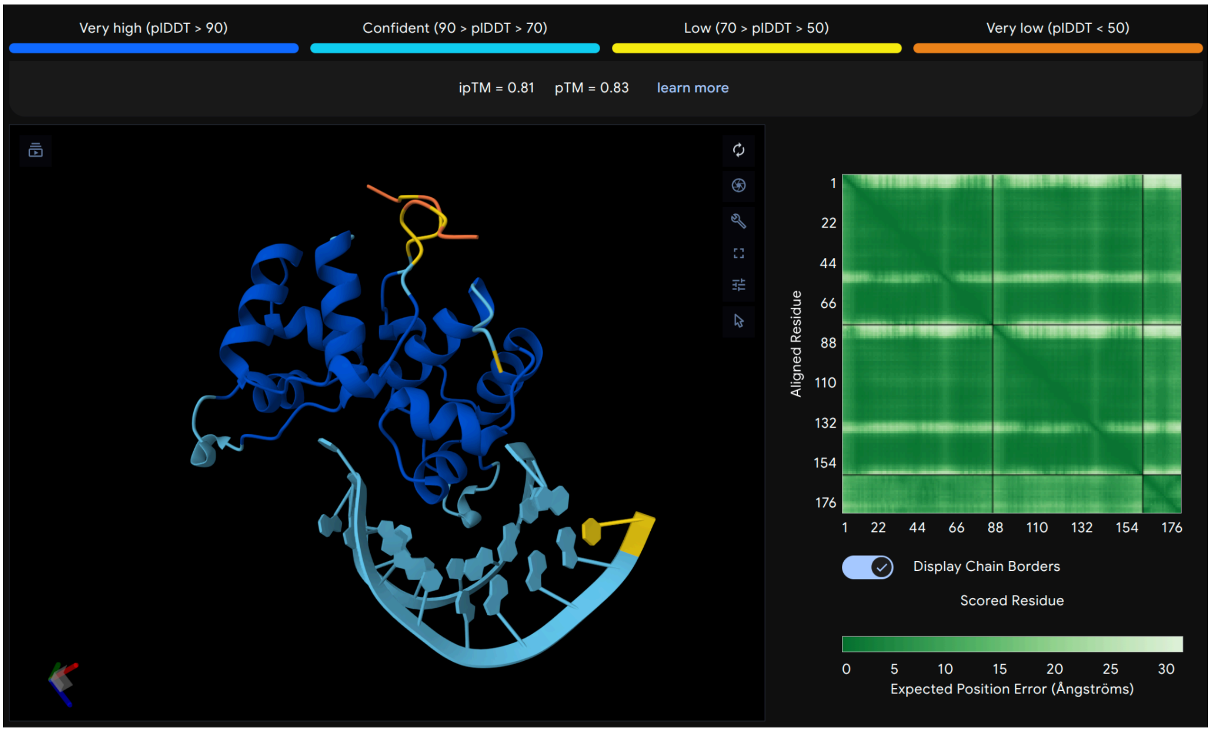


**Figure S4: AlphaFold3 prediction of Aca1 binding its predicted RNA motif.** The AlphaFold3 webserver was used for the prediction and visualization of the results (https://alphafoldserver.com/, Accessed 25.06.2026). For input sequence see **Supplementary Table S3**.

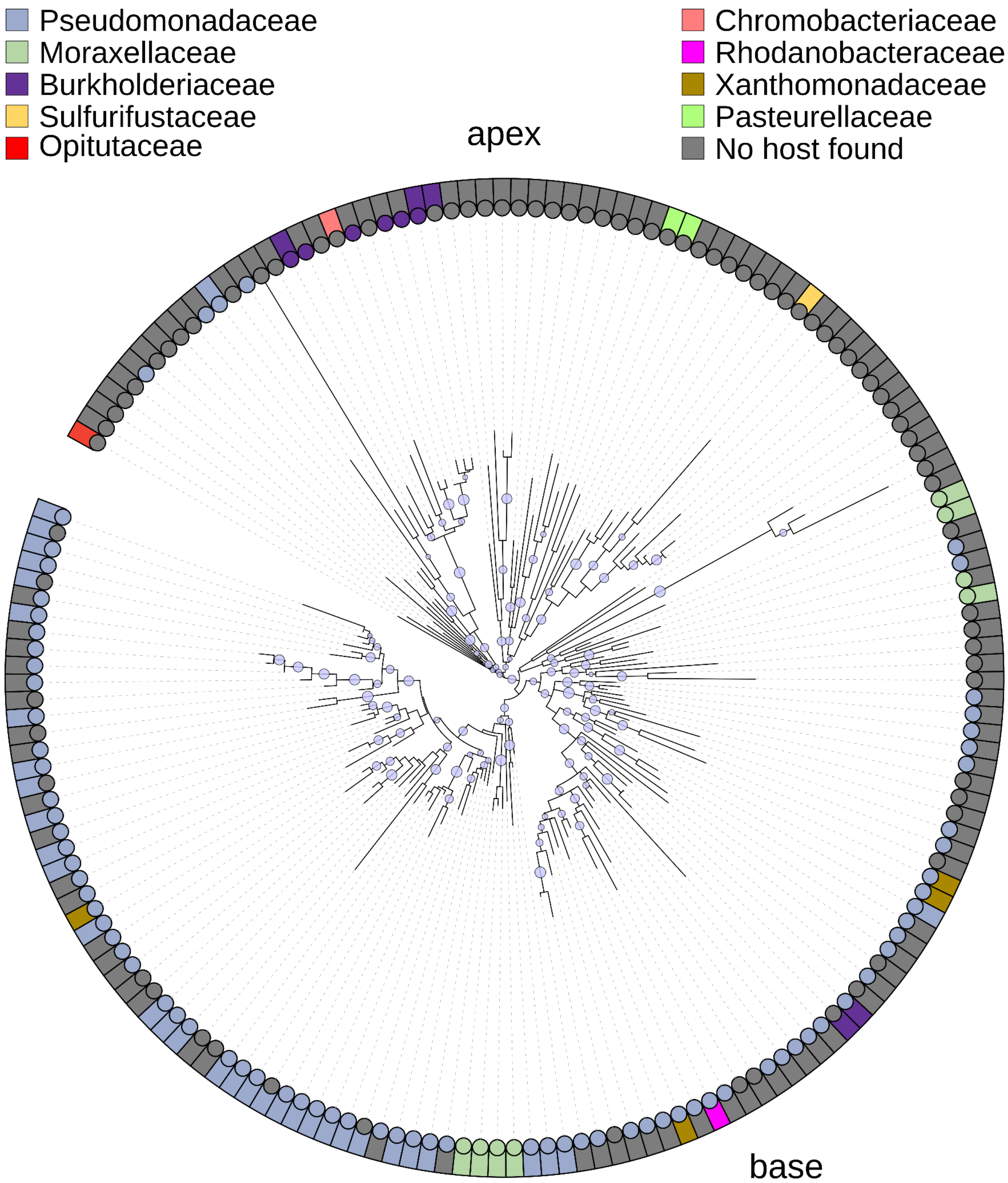


**Figure S5: Phylogenetic tree of Aca1 with bootstrap values and branch length.** This tree is the enlarged version of Figure 3H. Light blue circles indicate the bootstrap value with the smallest circle representing a bootstrap value of 0.7 and the largest 1.

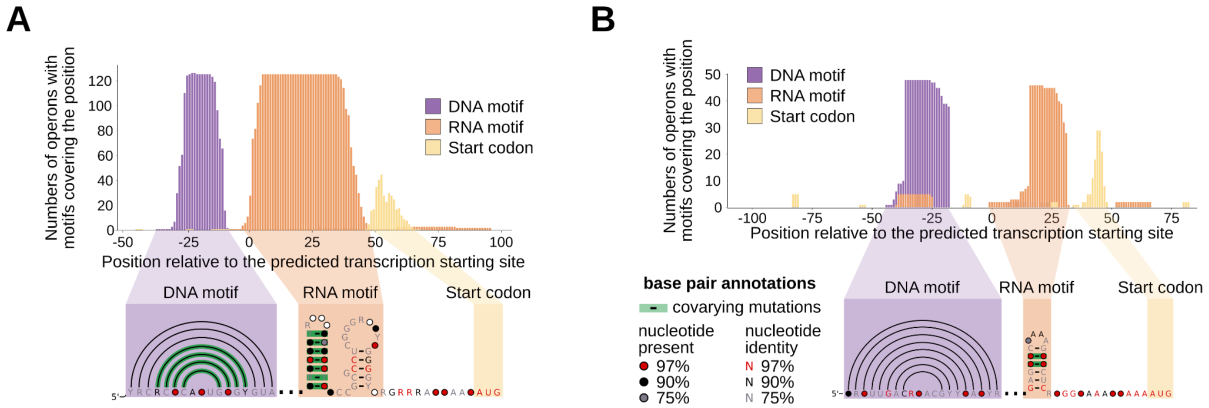


**Figure S6: Potential RNA motifs from Aca10 and Aca12 show sequence and positional conservation and covariation.** Position plot from the 5′ UTR of **(A)** *aca10* operons or **(B)** *aca12* operons, indicating the position of each found RNA motif (orange), upstream DNA motif (blue), and downstream start codon (yellow) relative to the predicted transcription starting site. Consensus sequence and RNA secondary structure of the conserved DNA binding site (blue) and the predicted RNA motif (orange) from an alignment of **(A)** 139 *aca10* operons or **(B)** 66 *aca12* operons. Nucleotide colour area as in Figure 2C.

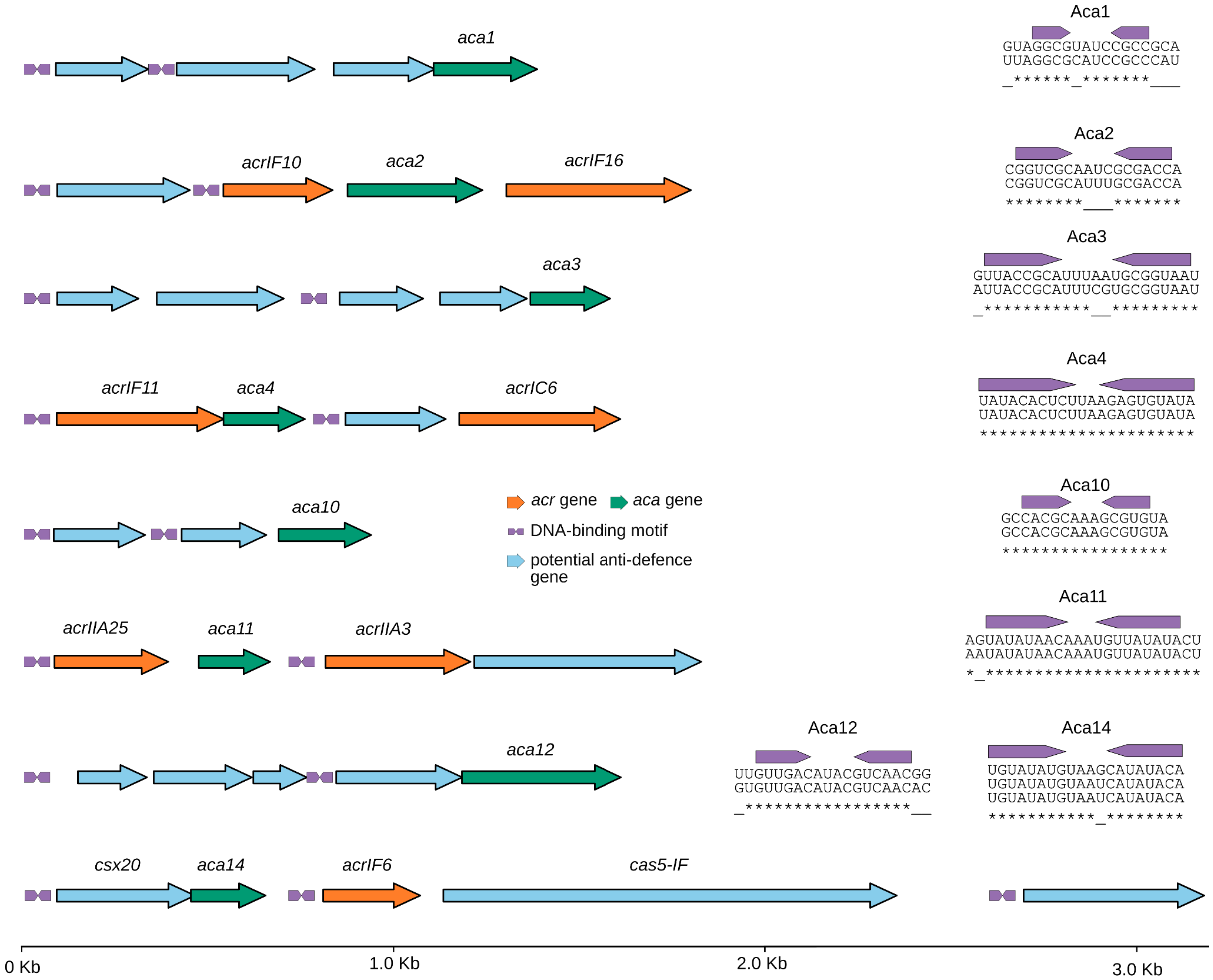


**Figure S7: Predicted multi-operon control by Aca proteins mediated by highly similar inverted-repeat DNA motifs. (Left)**, representative genomic arrangements showing predicted DNA-binding motifs upstream of up to three anti-defense operons linked to a single nearby aca gene. **(Right)**, sequence alignment of the corresponding predicted DNA-binding motifs from the operons shown on the left. Plum arrowheads indicate the inverted repeats; underlined positions are non-conserved, and asterisks indicate conserved nucleotides. Nucleotide identifiers: Ga0451703_0001175:1300-2750, JGVI01000034:98130-99925, Ga0099518_100418:26500-28100, Ga0198845_11:20000-21525, Ga0453721_0006603:6500-7475, AGZY01000007:268300-266400, Ga0099615_1000167:10590-12150, and DAOHFP010000061:1-3250.

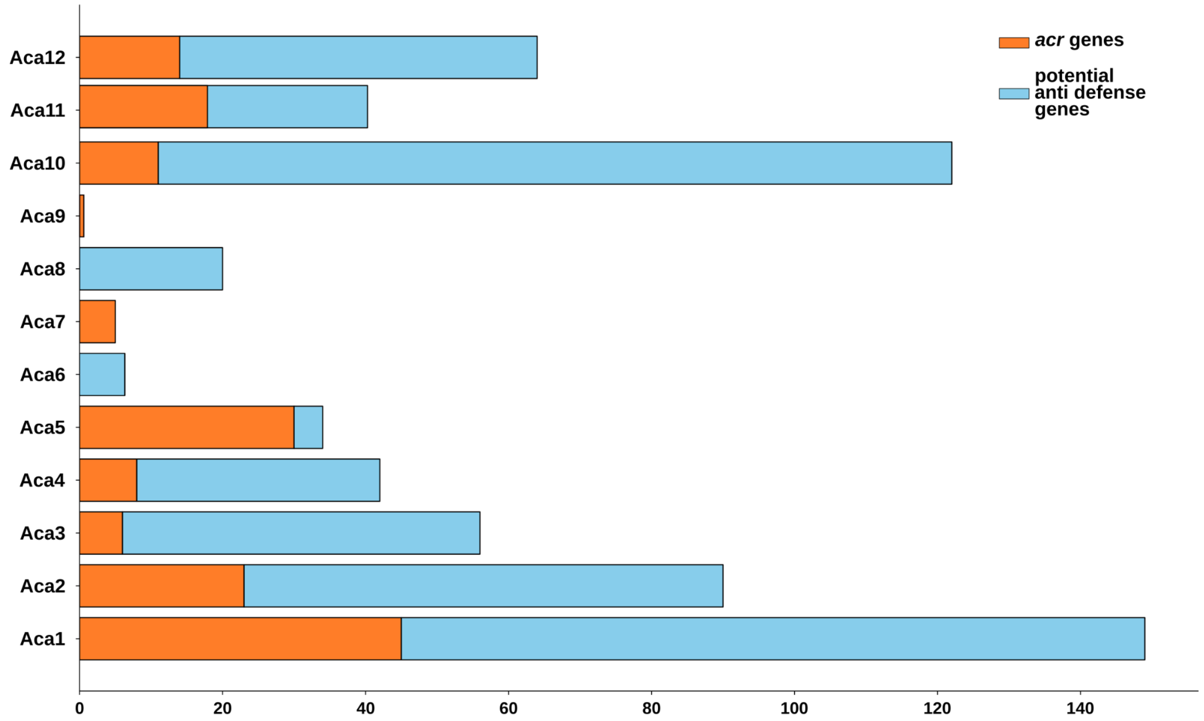


**Figure S8: Annotation of the proteins directly downstream of the predicted Aca DNA motifs.** On the left (orange) are annotated *acr* genes identified based on the dbAPIS database, while on the right (blue) are genes of unknown function but with potential anti-defense roles. Only *acr-aca* operons in the IMG/VR database[3] were analysed.

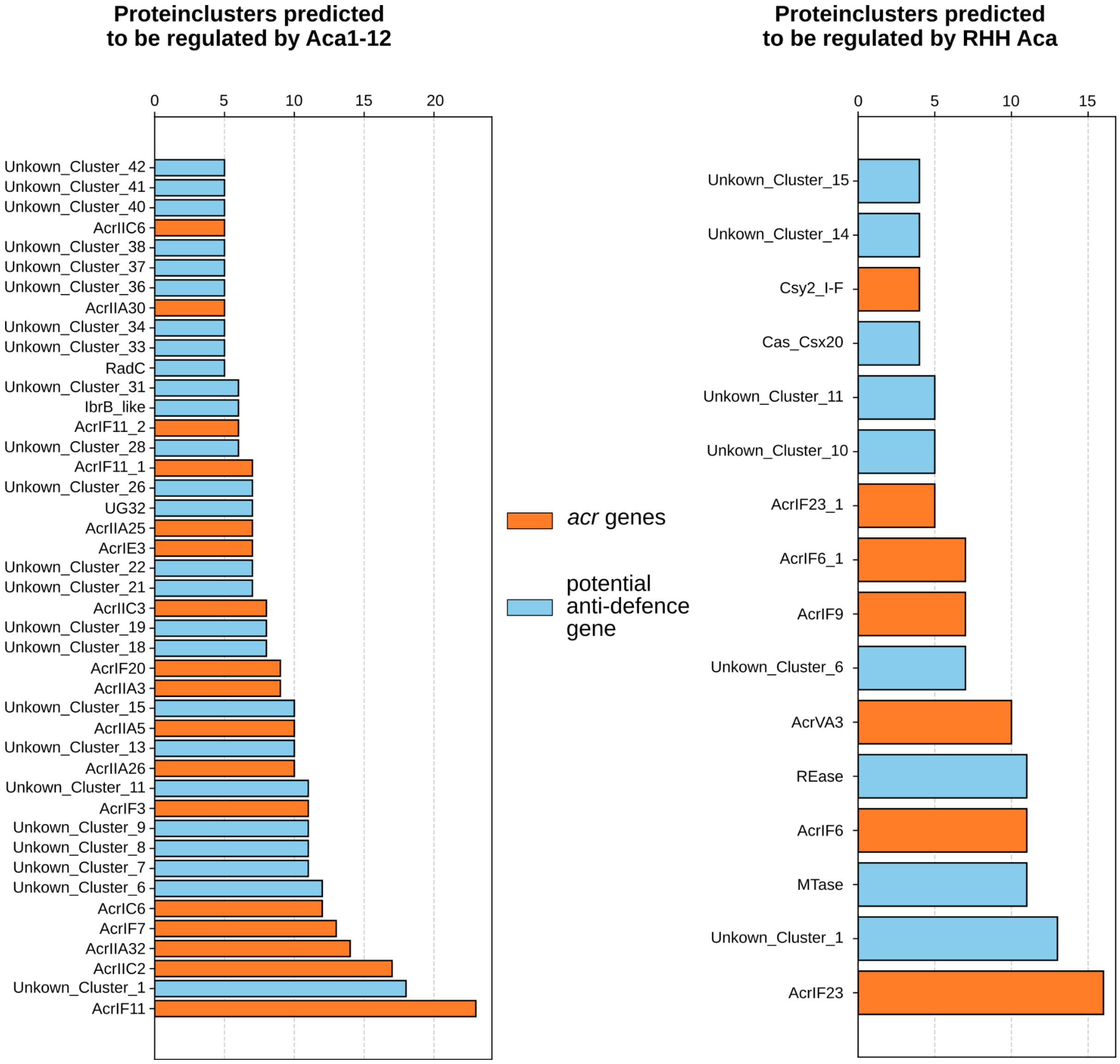


**Figure S9: Protein predicted to be regulated by Aca1-12 and Aca14 (RHH).** Protein clusters were annotated with DefenseFinder HMMs and the CDD database. Homologs of verified *acr* genes are orange and potential anti-defense genes are light blue **(Left)** Over 40 protein clusters with potential (n=25) or verified (n=17) anti defense function are predicted to be regulated by Aca1 to Aca12. Clusters with less than 5 proteins were not plotted. **(Right)** 15 protein clusters with potential (n=9) or verified (n=6) anti defense functions are predicted to be regulated by RHH Aca. Clusters with less than 4 proteins were not plotted.

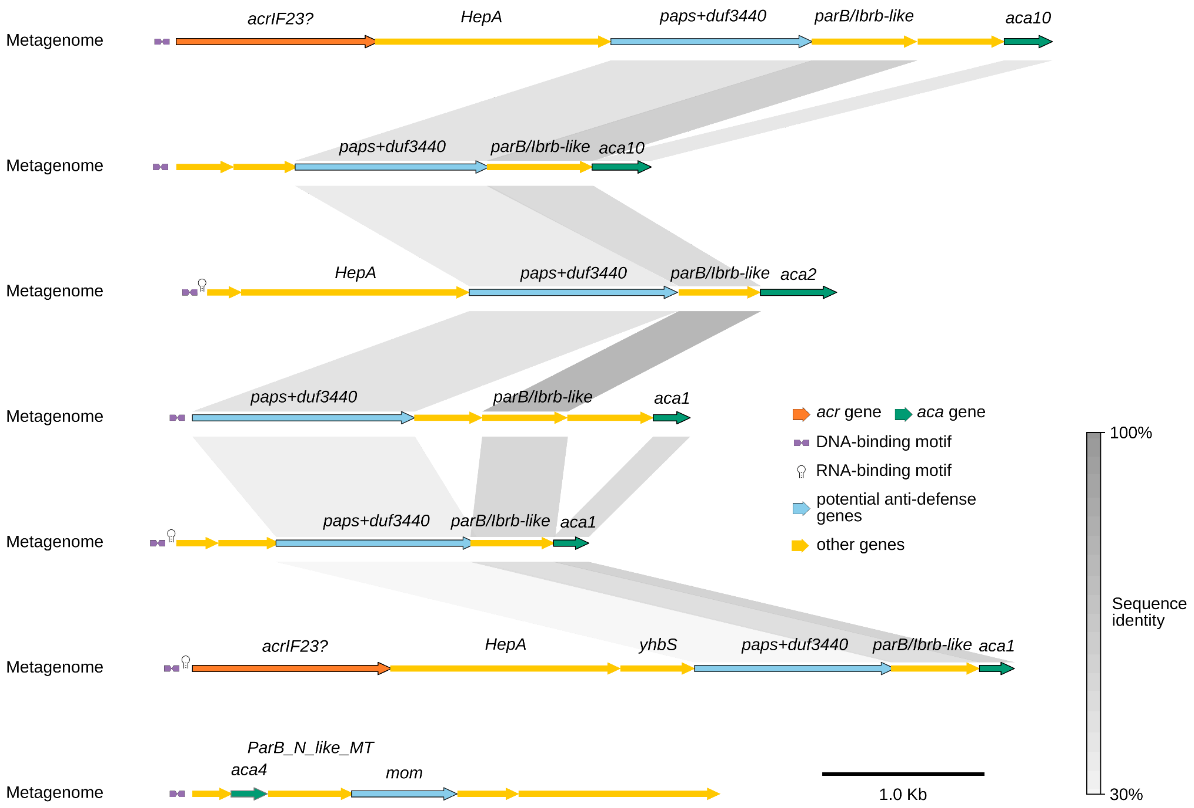


**Figure S10: Synteny plot of Phosphoadenosine phosphosulphate reductase and *mom* genes regulated by Aca proteins.** Operons were found in the IMG/VR database[3] and annotated with the CDD database[4] and DefenseFinder HMMs[5]. Nucleotide identifiers: Ga0074422_100915: 13430..20800, Ga0114347_1002298: 10000..13000, Ga0315268_10000735: 12000..15500, Ga0364483_10237: 42300..45800, Ga0376445_001921: 2500..5000, Ga0451687_0000891: 5200..10300, and Ga0453122_0006624: 5100..10100.

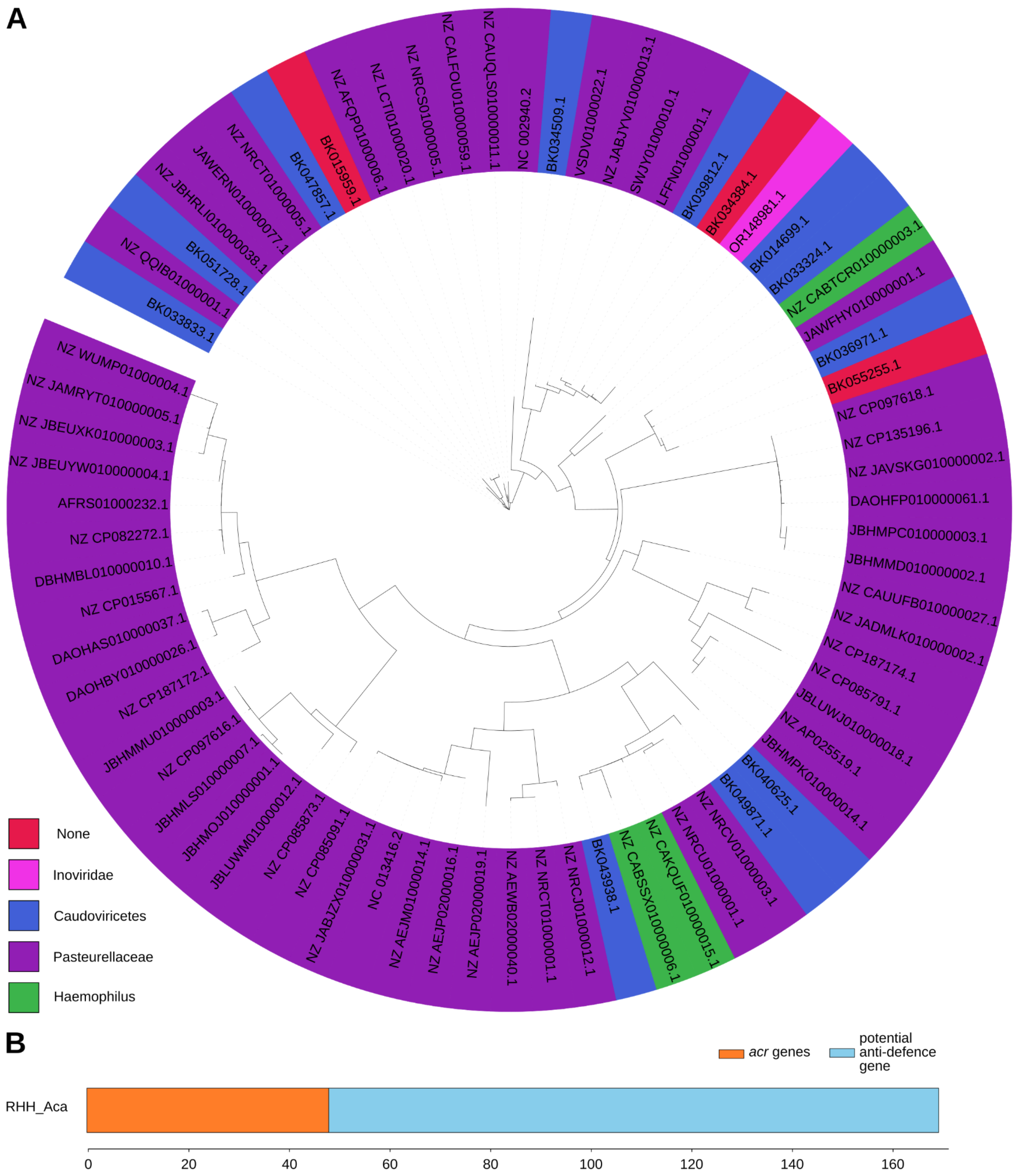


**Figure S11: Biological entities with the novel RHH Aca protein. (A)** Unrooted phylogenetic tree of RHH domain proteins with predicted regulatory sequences. **(B)** Genes in operons predicted to be regulated by the RHH-domain Aca protein. On the left (orange) are annotated *acr* genes identified via the dbAPIS database; on the right (blue) are genes of unknown function with putative anti-defence roles.

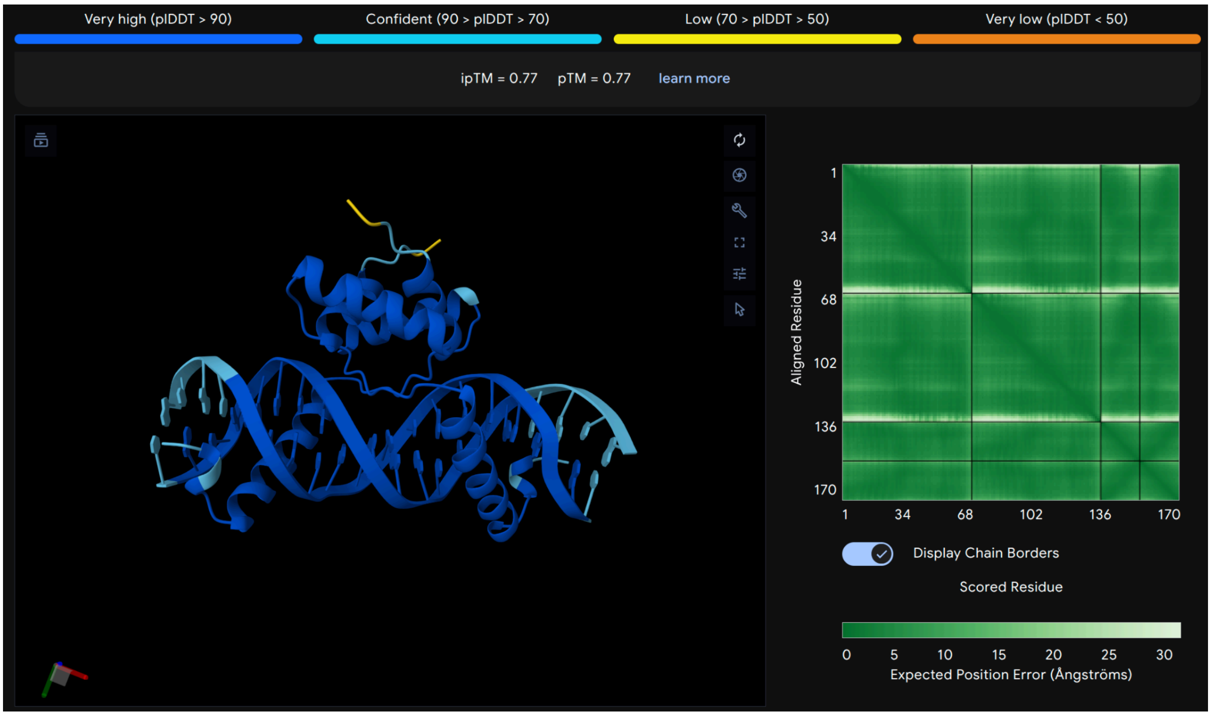


**Figure S12: AlphaFold3 prediction of RHH-domain protein binding its predicted DNA inverted repeat.** The AlphaFold3 webserver was used for the prediction and visualization of the results (https://alphafoldserver.com/, Accessed 25.06.2026). For input sequences, see **Supplementary Table S1**.

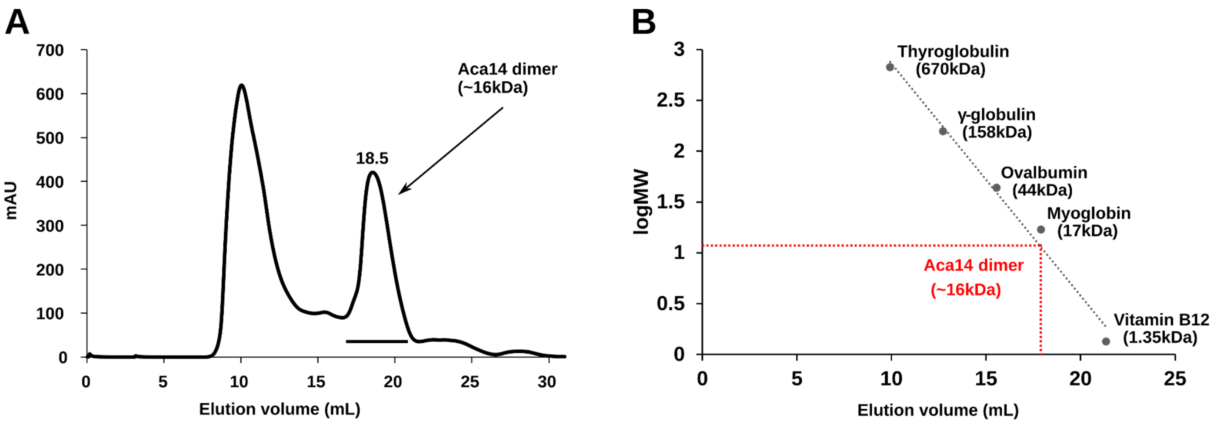


**Figure S13: Size exclusion chromatography of Aca14. (A)** Elution profile of Aca14 with the peak of the Aca14 dimer at 18.5 ml. **(B) SEC** calibration curve (black dashed line) using the indicated proteins of different molecular weights. The elution volume of the Aca14 dimer and its corresponding molecular weight are indicated by the red dotted line.

**Table S1. Permutation-based parsimony analysis of RNA-cluster and host-family clustering on Aca1 phylogenies.** Summary of tree–trait association tests performed on fixed Aca1 amino-acid phylogenies generated from unclustered sequences or sequences dereplicated at 95% and 90% amino-acid identity. RNA-cluster identity and predicted host family were treated as discrete tip traits, and phylogenetic clustering was quantified using the Fitch parsimony score with significance assessed by tip-label permutation. Lower observed parsimony scores indicate fewer inferred state changes and therefore stronger clustering of identical trait values on the tree. For the rna_cluster, ‘0’ indicates the Pseudomonadaceae, ‘1’ the Moraxellaceae, and ‘2’ the Burkholderiaceae cluster.

| **Metric** | **No clustering** | **95% identity clustering** | **90% identity clustering** |
|---|---:|---:|---:|
| RNA-cluster trait tested | rna_cluster | rna_cluster | rna_cluster |
| Number of tips | 190 | 111 | 93 |
| Number of states | 3 | 3 | 3 |
| State counts | 0: 154; 1: 30; 2: 6 | 0: 97; 1: 8; 2: 6 | 0: 79; 1: 8; 2: 6 |
| Observed parsimony score | 6 | 5 | 5 |
| Null mean parsimony score | 34.2320 | 13.7049 | 13.6630 |
| Null SD | 1.2944 | 0.5325 | 0.5732 |
| Z-score | -21.8109 | -16.3475 | -15.1141 |
| Empirical one-sided p-value | 0.000010 | 0.000010 | 0.000010 |
| **Host-family trait tested** | host_family | host_family | host_family |
| Number of tips | 126 | 56 | 44 |
| Number of states | 5 | 5 | 5 |
| State counts | Pseudomonadaceae: 89; Moraxellaceae: 29; Burkholderiaceae: 3; Xanthomonadaceae: 4; Rhodanobacteraceae: 1 | Pseudomonadaceae: 41; Moraxellaceae: 7; Xanthomonadaceae: 4; Burkholderiaceae: 3; Rhodanobacteraceae: 1 | Pseudomonadaceae: 29; Moraxellaceae: 7; Xanthomonadaceae: 4; Burkholderiaceae: 3; Rhodanobacteraceae: 1 |
| Observed parsimony score | 9 | 10 | 9 |
| Null mean | 34.2323 | 14.5233 | 14.3514 |

| parsimony score | | | |
|---|---|---|---|
| Null SD | 1.6114 | 0.6653 | 0.7769 |
| Z-score | -15.6583 | -6.7988 | -6.8884 |
| Empirical one-sided p-value | 0.000010 | 0.000060 | 0.000010 |
| Interpretation | significant phylogenetic clustering | significant phylogenetic clustering | significant phylogenetic clustering |

**Table S2: Sequences and stoichiometries used for AF3 to predict the DNA-protein interaction of the RHH-domain protein.**

| Stoichiometries | Amino acid/DNA sequences | ipTM (Stoichiometries) | pTM (Stoichiometries) |
|---|---|---|---|
| 1,2,4 | MRYENLSLERRRAISKNADAYNKENYKTITVRLLPETFEKLEQIIKKEGLSRNKAINKLIEEHKKP | 0.11 (1), 0.77 (2), 0.3 (4) | 0.4 (1), 0.77 (2), 0.41 (4) |
| 2 | TGTATATGTAAGCATATACA | | |

**Table S3: Sequences and stoichiometries used for AF3 to predict the DNA-protein and RNA-protein interaction of Aca1.**

| Stoichiometries | Sequence | ipTM (Stoichiometries) | pTM (Stoichiometries) |
|---|---|---|---|
| 2 (Aca1) | MRFPGVKTPDASNHDPDPRYLRGLLKKAGISQRRAAELLGLSDRVMRYYLSEDIKEGYRPAPYTVQFALECLANDPPSA | | |
| 1, 2 (RNA) | GCCAGCCACAACGGCGAGGC | 0.81 (1), 0.74 (2) | 0.83 (1), 0.79 (2) |

**Table S4: Plasmids used in this study.**

| Plasmid | Function | Reference or construction notes |
|---|---|---|
| pBAD30 | Empty vector for insertion of *aca14* under $P_{araBAD}$ control; $Amp^R$; pACYC184/p15A replicon | https://doi.org/10.1128/jb.177.14.4121-4130.1995 |
| pPF1439 | Promoterless *eyfp* reporter plasmid; $Cm^R$; pBR322 replicon | https://doi.org/10.1038/s41564-020-00822-7 |
| pPF4545 | pBAD30-derived expression plasmid | this study; insertion of |

| | | |
|---|---|---|
| | for *aca14* from *Pasteurella multocida* P2095 | gBlock PF10050 into pBAD30 via EcoRI+HindIII |
| pPF4696 | pPF1439-derived transcriptional fusion of *aca14*-controlled upstream promoter (operon 1) with an unrelated 5′ UTR and *eyfp* | this study; insertion of gBlock PF10102 into pPF1439 via SpeI+NsiI |
| pPF4697 | like pPF4696 but with mutated inverted repeats | this study; insertion of gBlock PF10103 into pPF1439 via SpeI+NsiI |
| pPF4698 | pPF1439-derived transcriptional fusion of *aca14*-controlled downstream promoter (operon 2) with an unrelated 5′ UTR and *eyfp* | this study; insertion of gBlock PF10104 into pPF1439 via SpeI+NsiI |
| pPF4699 | like pPF4698 but with mutated inverted repeats | this study; insertion of gBlock PF10105 into pPF1439 via SpeI+NsiI |
| pET28a-Aca1 | Expression plasmid encoding N-terminal His-tagged Aca1 from *Pseudomonas aeruginosa* phage JBD30; Kan$^R$; f1 replicon | this study; synthesized and cloned into pET28a by Bionics |
| pET28a-Aca12 | Expression plasmid encoding N-terminal His-tagged Aca12 from bacteriophage sp.; Kan$^R$; f1 replicon | this study; synthesized and cloned into pET28a by Bionics |
| pET28a-Aca14 | Expression plasmid encoding N-terminal His-tagged Aca14 from *Pasteurella multocida*; Kan$^R$; f1 replicon | this study; synthesized and cloned into pET28a by Twist Bioscience |

**Table S5: DNA oligonucleotides used in this study.**

| **Name** | **Sequence** |
|---|---|
| PF10050 | TCGTCTTCACCTCGAGAAATCGAATTCAGGAGGACACGGATGCGGTATGAAAATTTATCACTAGAACGAAGACGGGCAATCAGCAAAAATGCTGATGCTTATAATAAAGAAAATTACAAGACGATAACTGTCAGGCTATTGCCTGAAACTTTCGAAAAGCTAGAACAGATAATCAAGAAAGAAGGCTTGTCGAGAAACAAAGCAATAAACAAGCTCATTGAAGAATACAAAAAGCCCTAAAAGCTTCCTGTTGATAGATCCAGTAATGAC |
| PF10102 | TCGTCTTCACCTCGAGAAATCACTAGTAAAATAAAAAATACTTGATTTTTTATCCTGTATATGAAATCATATACAAAACCCATCTTAGTATATTAGTTAAGTATAAGAAGGAGATATACATATGCATCCTGTTGATAGATCCAGTAATGAC |
| PF10103 | TCGTCTTCACCTCGAGAAATCACTAGTAAAATAAAAAATACTTGATTTTTTATCCTGTATATGAAATCATAAAGAAAACCCATCTTAGTATATTAGTTAAGTATAAGAAGGAGATATACATATGCATCCTGTTGATAGATCCAGTAAT |

| | |
|---|---|
| | GAC |
| PF10104 | TCGTCTTCACCTCGAGAAATCACTAGTTAGATAAAAAATCTTGACTATTTAAAATGTATATGTAATCATATACACAACCCATCTTAGTATATTAGTTAAGTATAAGAAGGAGATATACATATGCATCCTGTTGATAGATCCAGTAATGAC |
| PF10105 | TCGTCTTCACCTCGAGAAATCACTAGTTAGATAAAAAATCTTGACTATTTAAAATGTATATGTAATCATAAAGACAACCCATCTTAGTATATTAGTTAAGTATAAGAAGGAGATATACATATGCATCCTGTTGATAGATCCAGTAATGAC |
| aca12_sense | FAM-AAAAATACTTGACGAACATCAAGTAATATG |
| aca12_antisense | CATATTACTTGATGTTCGTCAAGTATTTTT |
| aca14_IRup_sense | FAM-TTGATTTTTTATCCTGTATATGAAATCATATACAAAT |
| aca14_IRup_antisense | ATTTGTATATGATTTCATATACAGGATAAAAAATCAA |
| aca14_IRup_unlabeled | same as aca14_IRup_sense without the FAM label |
| aca14_IRdown_sense | FAM-TTGACTATTTAAAATGTATATGTAATCATATACACAG |
| aca14_IRdown_antisense | CTGTGTATATGATTACATATACATTTTAAATAGTCAA |
| aca14_IRdown_unlabeled | same as aca14_IRdown_sense without the FAM label |
| Nonspecific_sense | TTCTAATACGACTCACTATAGTGCAACCTTCATTTCCCTGCGTTTTAGAGCTAGA |
| Nonspecific_antisense | TCTAGCTCTAAAACGCAGGGAAATGAAGGTTGCACTATAGTGAGTCGTATTAGAA |

**Table S6: RNA probes used for Aca1 EMSAs.** All probes are labelled with IRDye® 800CW at their 5′ ends.

| **Probe** | **Sequence** |
|---|---|
| *aca1* (PF8279) | AUGCCAGCCACAACGGCGAGGCGCCAACAAGGAUUCAAACCAUGAAGUUCAUCAAAUACC |
| *aca2* (PF4483) | AUCGGUUCGAGAUGGCUCGAAUCGCUCCUAACGAGGAUUCCACAAUGUCUACUGCUUACA |

| | |
|---|---|
| *aca10* (PF9678) | GGCAGGCGUAAACGCCGACCACCGCCCCGGCGGAACCGGGC UUCCUGAUAGGAGCAACAU |
| *aca12* (PF9679) | AAGAUAAAGGAAGGGCCGAAAAGUUCGGCGGGGUAAAUGAAA AUGAAAAAAUUUUAUAUC |
